# DeepCINAC: a deep-learning-based Python toolbox for inferring calcium imaging neuronal activity based on movie visualization

**DOI:** 10.1101/803726

**Authors:** Julien Denis, Robin F. Dard, Eleonora Quiroli, Rosa Cossart, Michel A. Picardo

## Abstract

Two-photon calcium imaging is now widely used to infer neuronal dynamics from changes in fluorescence of an indicator. However, state of the art computational tools are not optimized for the reliable detection of fluorescence transients from highly synchronous neurons located in densely packed regions such as the CA1 pyramidal layer of the hippocampus during early postnatal stages of development. Indeed, the latest analytical tools often lack proper benchmark measurements. To meet this challenge, we first developed a graphical user interface allowing for a precise manual detection of all calcium transients from imaged neurons based on the visualization of the calcium imaging movie. Then, we analyzed the movies using a convolutional neural network with an attention process and a bidirectional long-short term memory network. This method is able to reach human performance and offers a better F1 score (harmonic mean of sensitivity and precision) than CaImAn to infer neural activity in the developing CA1 without any user intervention. It also enables automatically identifying activity originating from GABAergic neurons. Overall, DeepCINAC offers a simple, fast and flexible open-source toolbox for processing a wide variety of calcium imaging datasets while providing the tools to evaluate its performance.

**Significance statement:** Inferring neuronal activity from calcium imaging data remains a challenge due to the difficulty in obtaining a ground truth using patch clamp recordings and the problem of finding optimal tuning parameters of inference algorithms. DeepCINAC offers a flexible, fast and easy-to-use toolbox to infer neuronal activity from any kind of calcium imaging dataset through visual inspection.

## 1. Introduction

*In vivo* calcium imaging is widely used to study activity in neuronal microcircuits. Advances in imaging now allows for the simultaneous recording of several thousands of neurons (Stringer et al., 2019). One difficulty resides in how to infer single neuron activation dynamics from changes in fluorescence of a calcium indicator. A challenge is therefore to offer an analytical tool that would be scalable to the wide variety of calcium imaging datasets while providing reliable analysis.

State of the art computational tools to infer neuronal activity (such as CaImAn (Giovannucci et al., 2019; Pnevmatikakis et al., 2016)) are based on the deconvolution and demixing of fluorescence traces from segmented cells. However, in order to optimize the deconvolution parameters, a ground truth based on simultaneous targeted patch-clamp recordings and two-photon imaging is necessary (Chen et al., 2013; Evans et al., 2019).

Moreover, an analysis based on the fluorescence traces even after a demixing process can still be biased by overlapping cells (Gauthier et al., 2018). In a recent study from Gauthier and collaborators (Gauthier et al., 2018) analyzing calcium imaging data recorded in the region CA1 in adult rodents (Gauthier and Tank, 2018), 66% of the cells were reported as having at least one false transient and overall, among 33090 transients (from 1325 sources), 67% were considered as true, 13% as false and 20% were unclassified. Those contaminations increase the risk of misinterpretation of the data. Inferring neuronal activity from the developing hippocampus *in-vivo* is even more challenging due to several factors: 1-recurring network synchronizations are a hallmark of developing neuronal networks (Ben-Ari et al., 1989; Galli and Maffei, 1988; O’Donovan, 1989; Provine, 1972), which results in frequent cell co-activations, 2-the somata of pyramidal neurons are densely packed which results in spatial overlap, 3-Different calcium kinetics are observed in the same field of view (due to different cell types and different stages of neuronal maturation (Allene et al., 2012)). All these points are illustrated in Videos 1 and 2 (Region CA1 of the hippocampus from mouse pups). In addition, most methods do not offer solutions to evaluate the performance of neuronal activity inference on user datasets. To meet those challenges, we have developed a graphical user interface (GUI) that allows for such evaluation through data exploration and a method based on deep-learning to infer neuronal activity. Even if several deep-learning-based methods to infer neuronal activity from fluorescence signals have already been developed (Berens et al., 2018), none proposes a method directly based on raw two-photon imaging signals.

Action recognition from videos has seen recent important progress thanks to deep learning (Bin et al., 2018). Using a similar approach, we have trained a binary classifier on calcium imaging movies (allowing us to explore both the forward and backward temporal information among the whole sequence of video frames) to capture the fluorescence dynamics in the field of view and then predict the activity of all identified cells. It gave us the opportunity to take full advantage of the information contained in the movie in terms of dynamics and potential overlaps or other sources of contamination that might not be accessible when working only on fluorescence time courses.

To train the classifier a ground truth was needed. To our knowledge, no calcium imaging datasets from the developing hippocampus *in vivo* with simultaneous electrophysiological ground truth measurements are available. The most accurate ground truth would require targeted patch-clamp recordings with two-photon imaging on all the different hippocampal cell types with different calcium dynamics. This is technically difficult, time consuming and even more during development as the ground truth must be obtained from cells at various stages of maturation. As a result, we decided to base the ground truth on the visual inspection of raw movies using a custom-made GUI. It gives the advantages to work on any kind of calcium imaging dataset and to offer an easy tool to benchmark methods that infer neuronal activity.

The GUI offers a tool to precisely and manually detect all calcium transients (from onset to peak, which is the time when cells are active). We collected and combined a corpus of manual annotations from four human experts representing 37 hours of two-photon calcium imaging from 11 mouse pups aged between 5 to 16 postnatal days in the CA1 region using GCaMP6s. Almost 80 % of the labeled data was used to train the model, while the rest was kept to benchmark the performance. Then, movies were processed using a convolutional neural network with an attention mechanism and a bidirectional long-short term memory network (Hochreiter and Schmidhuber, 1997; LeCun and Bengio, 1995; Vaswani et al., 2017).

To evaluate the method, we used the ground truth as a benchmark. We found that this method reached human level performance and offered a better sensitivity and F1 score than CaImAn to infer neuronal activity in the developing hippocampus without any user intervention. Overall, DeepCINAC (Calcium Imaging Neuronal Activity Classifier) offers a simple, ergonomic, fast and flexible open-source toolbox for processing a wide variety of calcium imaging data while providing the tools to evaluate its performance.

## 2. Methods

In this section, we will describe all the necessary steps to build a deep learning neural network “DeepCINAC”. This toolbox was developed to analyze in vivo two-photon calcium imaging data acquired in the developing hippocampus (See § **<u>Experimental procedure and data acquisition</u>**). As a first step, we needed to set a ground truth that was established on the visualization of the recorded movie by three to four human experts (§ **<u>Ground truth</u>**). Then data are pre-processed (§ **<u>Data pre-processing and feature engineering and model description</u>**) and used to train the network (§ **<u>Computational performance</u>**). As a final step, we used labelled data to evaluate the performance of DeepCINAC (§ **<u>Performance evaluation</u>**). Tutorials and the source code are freely available online (§ **<u>Toolbox and data availability</u>**).

### 2.1 Experimental procedure and data acquisition

All experiments were performed under the guidelines of the French National Ethic Committee for Sciences and Health report on “Ethical Principles for Animal Experimentation” in agreement with the European Community Directive 86/609/EEC.

#### Viral injection

To induce widespread, rapid and stable expression of the calcium indicator GCaMP6s in hippocampal neurons at early postnatal stages, we intraventricularly injected a viral solution (pAAV.Syn.GCaMP6s.WPRE.SV40, Addgene #100843-AAV1) at P0 in mouse pups (Figure 1A-B). This injection protocol was adapted from already published methods (Kim et al., 2014, 2013). Mouse pups were anesthetized on ice for 3 to 4 minutes and 2 µL of the viral solution were injected in the left lateral ventricle which coordinates were estimated at the ⅖ of the imaginary line between the lambda and the eye at a depth of 400 µm. Expression of GCaMP was checked on slices and was sufficient for *in vivo* imaging as early as P5, which is consistent with already published data (Kim et al., 2014). In addition, GCaMP expression, brightness and kinetics of the reporter was then stable throughout all developmental stages used (data not shown).

**Figure 1:**
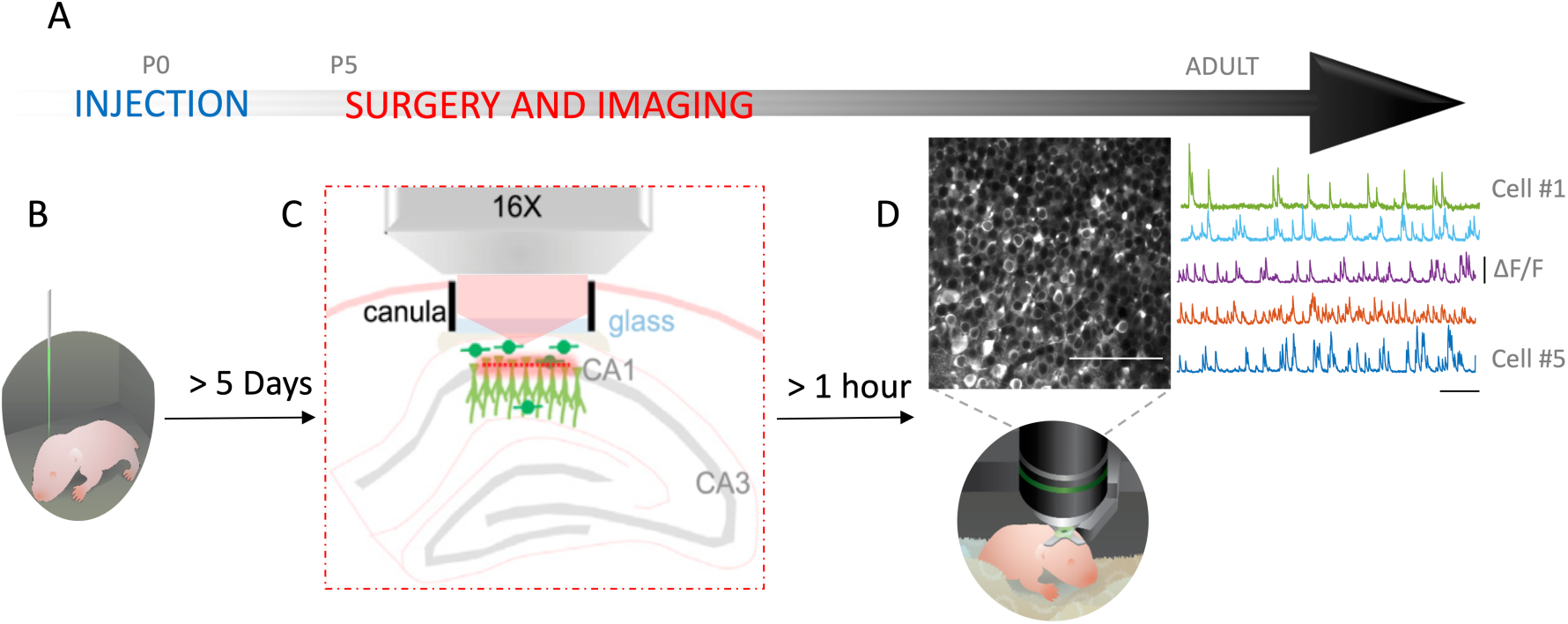
Experimental paradigm. ***1A***: *Experimental timeline*. ***1B***: *Intraventricular injection of GCaMP6s on pups (drawing) done at P0*. ***1C***: *Schematic representing the cranial window surgery*. ***1D***: *Top left: Imaged field of view. Scale bar: 100 µm. Top right: Activity of 5 random neurons in the field of view (variation of fluorescence is expressed as* Δf/f*). Scale bar 50 s. Bottom: Drawing of a head fixed pup under the microscope*.

#### Surgery

The surgery to implant a 3 mm large cranial window above corpus callosum was adapted from described methods (Dombeck et al., 2010; Villette et al., 2015). Anesthesia was induced using 3% isoflurane in a mix of 90% O2 - 10% air and maintained during the whole surgery (approximately 1:30h) between 1% and 2.5% isoflurane. Body temperature was controlled and maintained at 36°C. Analgesia was controlled using Buprenorphine (0.025mg/kg). Coordinates of the window-implant were estimated by eyes. The skull was removed and the cortex was gently aspirated until the external capsule / alveus that appears as a plexus of fibers was visible. Surface of the corpus callosum was protected with QuickSil (WPI) then the cannula with the window was implanted and fixed to the heaplate of the animal.

#### Imaging

Two-photon calcium imaging experiments were performed on the day of the surgery (Figure 1C-D) at least one hour after the end of the surgery. 12500-frames-long image series from a 400×400 μm field of view with a resolution of 200×200 pixels were acquired at a frame rate of 10.6 Hz (Figure 1D). We then motion-corrected the acquired images by finding the center of mass of the correlations across frames relative to a set of reference frames (Miri et al., 2011).

#### Cell segmentation

To detect cell contours, we used either the segmentation method implemented in suite2p (Pachitariu et al., 2017) or the Constrained Nonnegative Matrix Factorization (CNMF) implemented in CaImAn.

#### Activity inference

To infer activity we used the Markov chain Monte Carlo (MCMC) implemented in CaImAn on cell contours obtained from the CNMF of the toolbox. The MCMC spike inference was done as described (Pnevmatikakis et al., 2016). We used DeepCINAC predictions on both contours from suite2p and CaImAn.

### 2.2 Data visualisation: GUI

To visualize our data and explore the results from any spike inference method, we designed a graphical user interface (GUI) that provides a visual inspection of each cell’s activity (Figure 2). The GUI offers a set of functionalities allowing visualization of i) calcium imaging movies centered and zoomed on the cell of interest during a time window that includes a given transient, ii) sources, transient profiles and their correlations (as developed by Gauthier and collaborators), iii) transient fluorescence signal shape.

**Figure 2:**
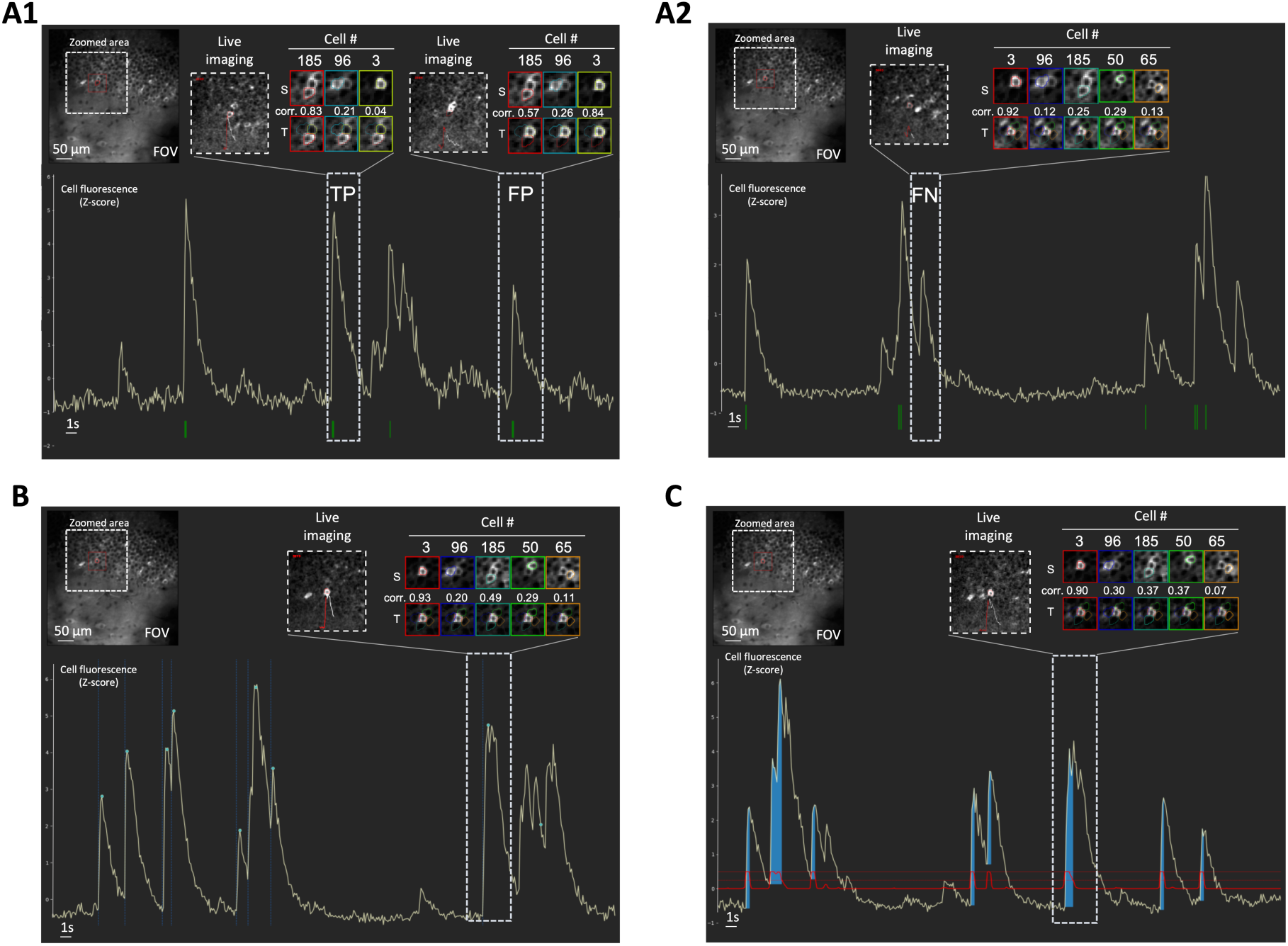
Examples of different uses of the GUI. The GUI can be used for data exploration (2A1-A2), to establish the ‘Ground Truth’ (2B) and to evaluate DeepCINAC predictions (2C). **2A:** The GUI can be used to explore the activity inference from any methods. The spikes inferred from CaImAn are represented by the green marks at the bottom. The GUI allows the user to play the movie at the time of the selected transient and visualize the transients and source profile of the cell of interest. **2A1:** Movie visualization and correlation between transient and source profiles allow the classification of the first selected transient as true positive (‘TP’) and the second selected transient as false positive (‘FP’). **2A2:** Movie visualization and correlation between transient and source profiles allow the classification of the selected transient as false negative (‘FN’). **2B:** The GUI can be used to establish a ‘Ground Truth’. In this condition it offers the user the possibility to manually annotate onset and peak of calcium transient. Onsets are represented by vertical dashed blue lines, peaks by green dots. **2C:** When the activity inference is done using DeepCINAC, the GUI allows the display of the classifier predictions. The prediction is represented by the red line. The dashed horizontal red line is a probability of one. The blue area represents time periods during which the probability is above a given threshold, in this example 0.5. Abbreviations: T: Transient profile, S: Source Profile, Corr: correlation, FOV: field of view

Additionally the GUI can be used to i) display the spike times from an inference method (Figure 2A1-2), ii) establish a ‘ground truth’ (Figure 2B), iii) visualize DeepCINAC predictions (Figure 2C).

The GUI was developed using Python and Tkinter package. It can read data from several formats including neurodata without borders files (Rübel et al., 2019; Teeters et al., 2015). More details on the GUI and a complete tutorial are available on GitLab (https://gitlab.com/cossartlab/deepcinac)

### 2.3 Ground truth

#### Activity inference

*Electrophysiological ground truth*. Ground truth data from experiments previously described were taken from crcns.org. (Chen et al., 2013; GENIE Project, 2015). Briefly, visual cortex neurons expressing the calcium indicator GCaMP6s were imaged while mice were presented with visual stimuli. 60 Hz two photon imaging and loose cell-attached recordings at 10 kHz were performed simultaneously. Using ImageJ software, we downsampled imaging data to 10 Hz by averaging every six frames and rescaled it to 1.2 µm/pixel. We considered a cell active during a rise time if a spike was detected during that time, and used the previously described GUI to convert those data in the cinac format so we could produce benchmarks and train a classifier using those data (see Table 1 and Table S1 for more details).

**Table 1:**
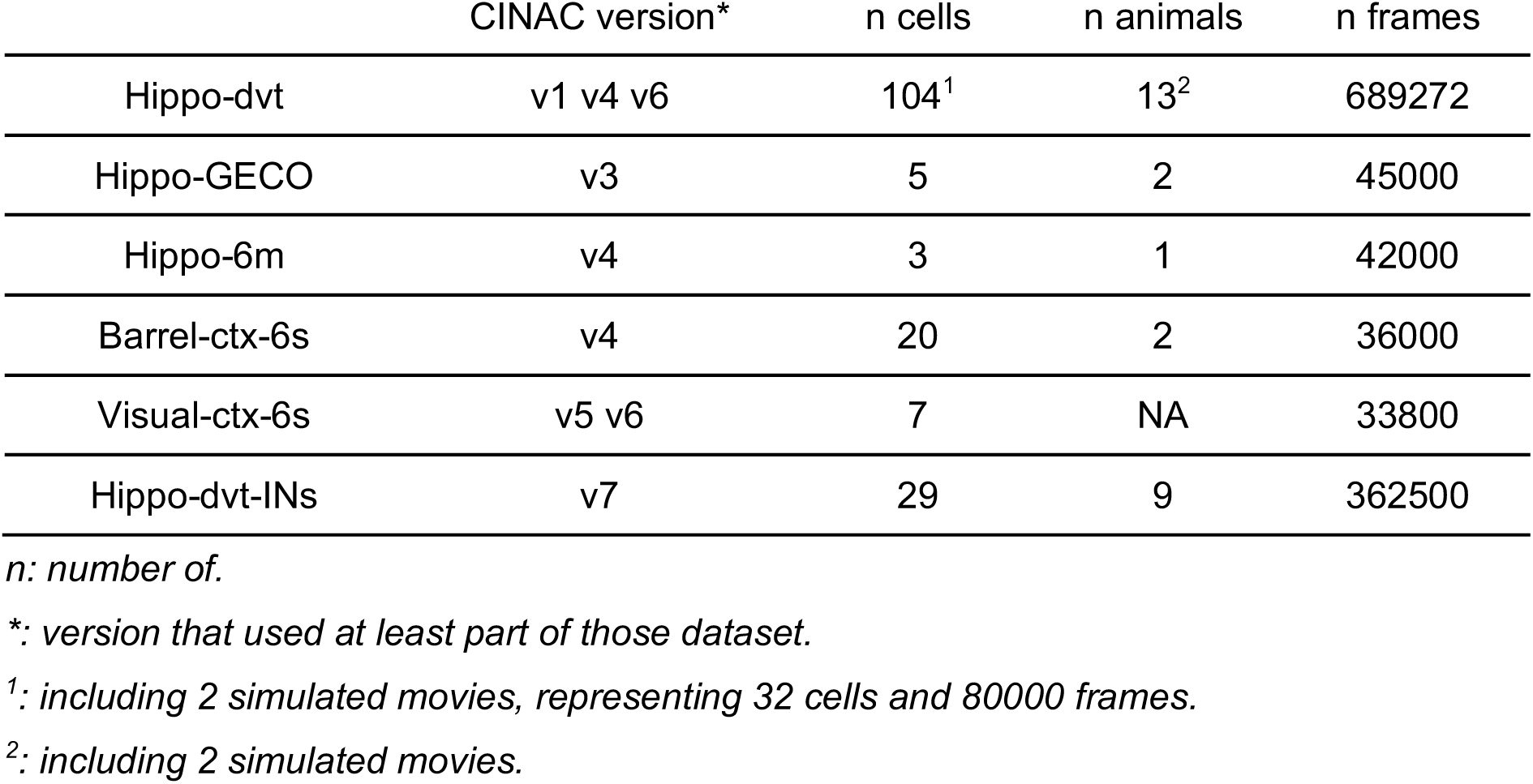
Data used to train the classifiers. Description of the datasets precising the number of frames, number of animals and field of views included, as well as the classifiers that used these datasets.

|  | CINAC version* | n cells | n animals | n frames |
| --- | --- | --- | --- | --- |
| Hippo-dvt | v1 v4 v6 | 104<sup>1</sup> | 13<sup>2</sup> | 689272 |
| Hippo-GECO | v3 | 5 | 2 | 45000 |
| Hippo-6m | v4 | 3 | 1 | 42000 |
| Barrel-ctx-6s | v4 | 20 | 2 | 36000 |
| Visual-ctx-6s | v5 v6 | 7 | NA | 33800 |
| Hippo-dvt-INs | v7 | 29 | 9 | 362500 |

*n*: number of.
\*: version that used at least part of those dataset.
<sup>1</sup>: including 2 simulated movies, representing 32 cells and 80000 frames.
<sup>2</sup>: including 2 simulated movies.

#### Activity inference

*Visual ground truth*. All functionalities of the GUI were used as criteria by each human expert to label the data. The ground truth was established based on two-photon calcium imaging from pups from 5 to 16 days old (see Table 1) in a four steps workflow as described in Figure 3. Data were selected and labeled at least by two independent human experts (Figure 3, step 1 & 2). We then combined those labels (Figure 3, step 3) and a final agreement was decided by three to four human experts (Figure 3, step 4). In addition, we trained another classifier for interneurons using transgenic pups in which only interneurons express the indicator (Melzer et al., 2012). As previously described, interneurons’ activity was labeled by three or four human experts and used to train an interneuron specific classifier (CINAC_v7, see Table 1). After training our classifier on a first set of cells, we used the predictions obtained on new data to establish additional ground truth based on the mistakes made on those data. At least two human experts labeled segments of 200 frames containing the wrong predictions. Additional visual ground truth was established by one human-expert (RD) on three other datasets from our lab using the GUI: i) GCAMP6s calcium imaging movies from the developing barrel cortex (‘Barrel-ctx-6s’, 1.5Hz, 1.2µm/pixel) (Modol et al., 2019), ii) GCaMP6m imaging movies (‘Hippo-6m’, 10 Hz, 2 µm/pixel) and iii) GECO imaging movies (‘Hippo-GECO’, 5 Hz, 2 µm/pixel) both from the adult hippocampus (see Table S1 for the details). For ‘Barrel-ctx-6s’ and ‘Hippo-GECO’, the CaImAn spike inference had already been performed by the original experimenter. We performed CaImAn spike inference on ‘Hippo-6m’.

**Figure 3:**
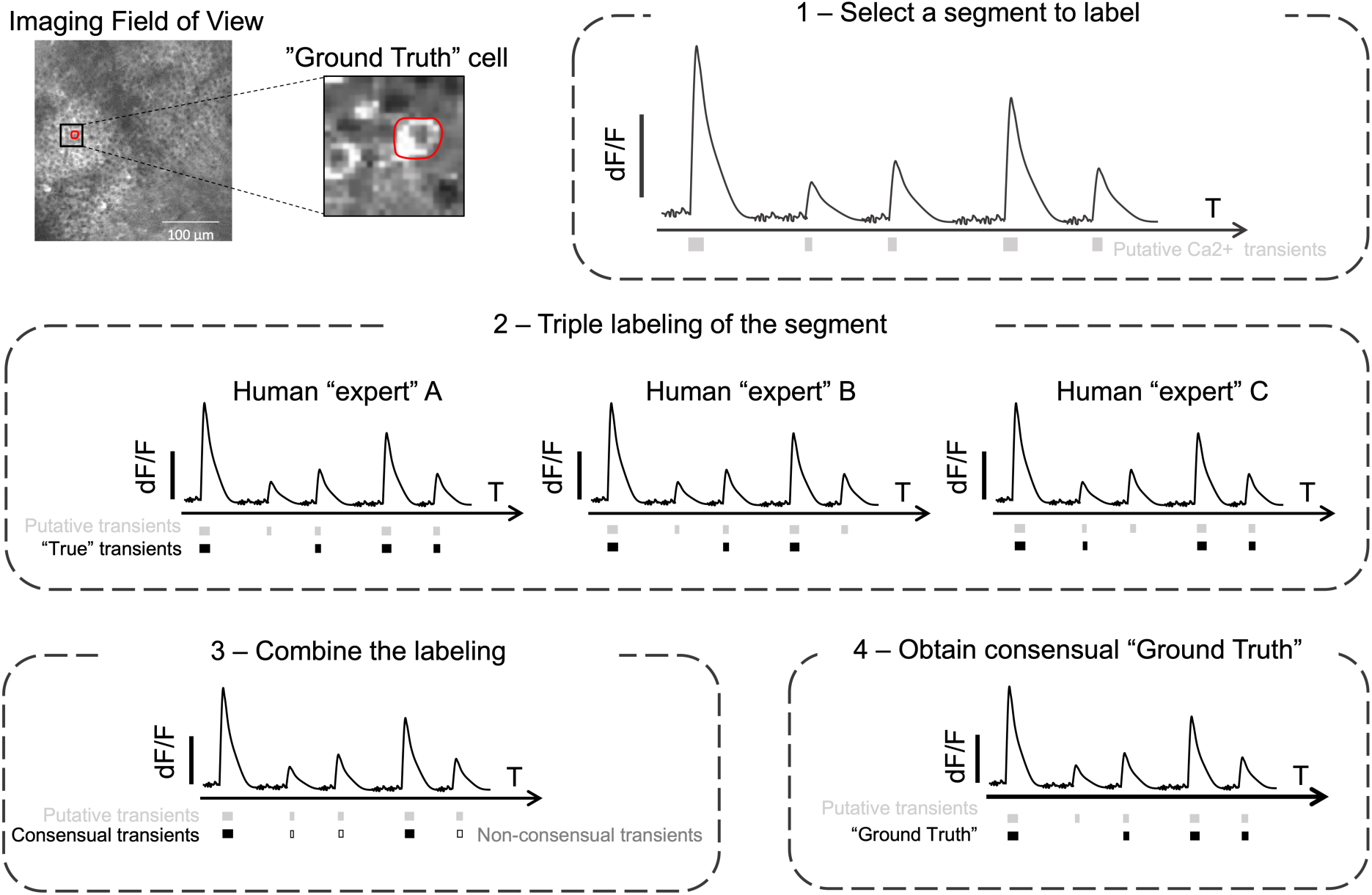
Workflow to establish the ‘Ground Truth’. First a cell was randomly chosen in the imaged field of view. **3-1** - All putative transients of the segment to label were identified for the onset to the peak of each calcium event. **3-2** - Three human experts (“expert” A, “expert” B, “expert” C) independently annotated the segment. Among all putative transients each human expert had to decide whether it was in his opinion a ‘true’ transient. **3-3** - The combination of the labelling lead to ‘consensual transients’ (i.e. ‘true’ transient for each human expert - black square) and to ‘non-consensual transients’ (i.e ‘true’ transient for at least one human expert but not all of them - open square). **3-4** - All ‘non-consensual transients’ were discussed and ‘ground truth’ was established.

#### Cell type ground truth

We used calcium imaging movies from GadCre (Melzer et al., 2012) positive animals injected with both h-SynGCaMP6s and Cre-dependent TdTomato to identify interneurons by the overlap of GCaMP6s and TdTomato signals. Using the GUI, we manually categorized 743 cells from 85 recordings among three categories: interneuron, pyramidal cell and noisy cell. 283 TdTomato expressing cells were categorized as interneurons. 296 cells were categorized as putative pyramidal cells based on their localization in the pyramidal layer, their shape and their activity. Finally 164 cells were categorized as noisy cells, determined by visually estimating their signal-to-noise ratio. We used a total of 643 cells (245 interneurons, 245 putative pyramidal cells and 153 noisy ones) to train the cell type classifier and 100 cells (38 interneurons, 51 putative pyramidal cells and 11 noisy ones, not included in the training dataset) were used to evaluate it.

### 2.4 Data pre-processing, feature engineering and model description

#### Data pre-processing and feature engineering

Calcium movies in tiff format were split into individual tiff frames to be efficiently loaded in real time during the data generation for each batch of data fed to the classifier. For any given cell, a batch was composed of a sequence of 100 frames of 25×25 pixels window centered on the cell body. The length of the batch was chosen to fit for interneurons activity (rise and decay time). The window size was adapted to capture the activity of cells overlapping the target cell. In a recording of 12500 frames, the number of transients ranges from 10 to 200 approximately. Thus the frames during which the cell is active (from onset to peak), represents a low percentage of the total data. Because manual labeling is time consuming, the data used as ground truth was limited in size. To overcome the issue of the imbalanced data and to enlarge the dataset, we used the following three approaches:

#### #1: Data augmentation (Perez and Wang, 2017)

temporal and spatial data augmentation were used. Temporal augmentation was used in that each block of 100 frames was overlapping with each other using a sliding window of 10 frames of length. Spatial augmentation took the form of transformations such as flip, rotation or translations of the images. The data augmentation was done online, meaning that the transformations were done on the mini-batches that the model was processing. This allowed avoiding memory consumption and generating a dataset on multiple cores in real time.

#### #2: Simulated data

To balance our dataset, and increase the ability of the network to predict a fake transient as false, we have simulated calcium imaging movies with a higher rate of overlapping activity than our dataset (an example of artificial movie is available online on the gitlab page, alongside the source code: https://gitlab.com/cossartlab/deepcinac). We started by collecting more than 2000 cell contours from several movies that were segmented using suite2p. We randomly picked contours to build a cell map, with sixteen cells for which one to four cells are overlapping it. We then generated for each cell an activity pattern, with a randomly chosen number of transients (from 2 to 16 for 1000 frames, 1.2 to 9.6 transients/min for a 10 Hz sampling rate) and duration of rise time (from 1 to 8 frames, 100 to 800 ms for a 10 Hz sampling rate), following a random distribution. To simulate the fluorescence signal, on the rise time we use a linear fit from the onset to peak, for the decay we use an exponential decay with a decay from 10 to 12 frames of duration. To generate the calcium imaging movie, we decided on a basal level of activity and then we adjusted the intensity of pixels in the cell for each frame according to the amplitude of the cell fluorescence, pixels in the cell have a different weight depending if they are in the soma or not, their intensity being lower in the nucleus. We finally added some gaussian noise on every frame.

#### #3: Data stratification

In order to balance the data, we used data augmentation on selected movie segments (underrepresented segments), and excluded others (overrepresented segments) from the training data set. After data stratification, we obtained approximately 60% of the movie segments containing at least one real transient, 30% at least one fake transient without real ones and 10% without transients. We were then able to be more precise over the proportion of segments with multiple transients or cropped transients. We gave higher weights to segments containing fake transients in order for the network to adjust the accuracy accordingly.

The data augmentation, simulated data and stratification were used only to produce the training dataset and not the validation dataset.

#### Model description

We designed a joint model combining a forward-pass long-short-term memory (LSTM), a backward-pass LSTM and convolutional neural network (CNN) features. In order for the bi-directional LSTM to focus on relevant information, we reinforced it by an attention process at the stage of encoding similar to previous work (Bin et al., 2018; Rémy, 2019). The model was designed using Python and Keras library (Chollet, 2015), see Figure 4).

**Figure 4:**
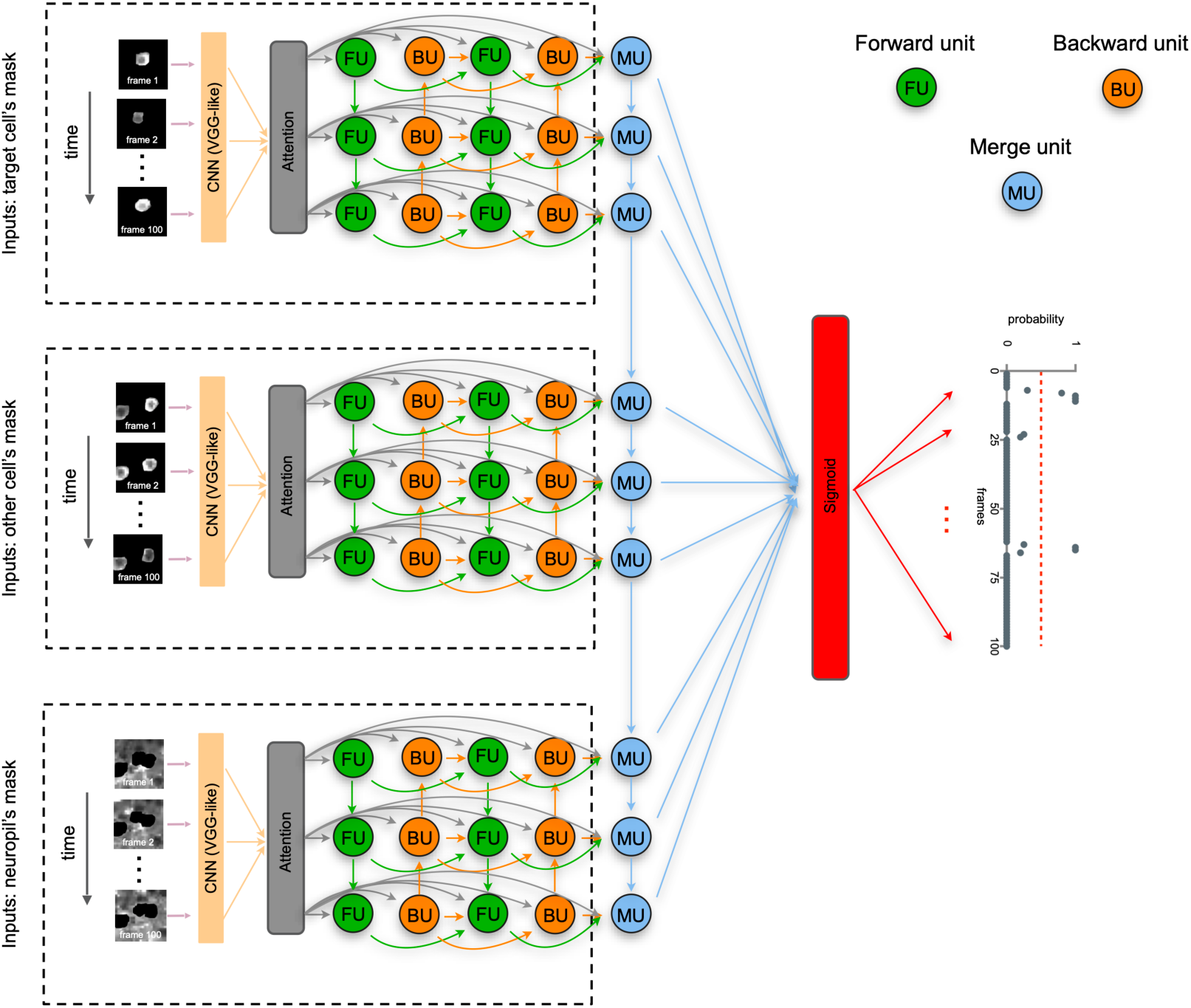
Architecture of DeepCINAC neural network. As a first step, for each set of inputs of the same cell, we extract CNNs features of video frames that we pass to an attention mechanism and feed the outputs into a forward pass network (FU, green units) and a backward pass network (BU, orange units), representing a bi-directionnal LSTM. Another bi-directionnal LSTM is fed from the attention mechanism and previous bi-directionnal LSTM outputs. A LSTM (MU, blue units) then integrate the outputs from the process of the three types of inputs to generate a final video representation. A sigmoid activation function is finally used to produce a probability for the cell to be active at each given frame given as input.

The model used to predict the cell activity takes three inputs, each representing the same sequence of 100 frames (around 10 seconds of activity). Each frame had dimensions of 25×25 pixels, centered around the cell of interest, whose activity we want to classify. The first input has all its pixels set to zero except for the mask of the cell of interest (cell activity). The second input has all its pixels set to zero except for the mask of the cells that intersect the cell of interest (overlapping activity). The final input has the cell of interest and the one intersecting its pixels set to zeros (neuropil activity). That way, the model has all the information necessary to learn to classify the cell’s activity according to its fluorescence variation.

The model used to predict the cell type takes two inputs, each representing the same sequence of 500 frames (around 50 seconds of activity). Each frame had dimensions of 20×20 pixels, centered around the cell of interest, whose cell type we want to classify. The first input has all its pixels set to zero except for the mask of the cell of interest (cell activity). The second input has all its pixels.

We used dropout (Srivastava et al., 2014) to avoid overfitting, but no batch normalization. The activation function was swish (Ramachandran et al., 2017). The loss function was binary cross-entropy and the optimizer was RMSprop. To classify cell activity the output of the model was a vector of length 100 with values between 0 and 1 representing the probability for the cell to be active at a given frame of the sequence. To classify the cell type (interneuron, pyramidal cell or noisy cell), the output was three values ranging from 0 to 1 and whose sum is equal to 1, representing the probability for a cell to be one of those three cell types.

### 2.5 Computational performance

#### Classifier training

We trained the classifier on a Linux-based HPC cluster where 4 CPUs (Intel(R) Xeon(R) CPU E5-2680 v3), 320 GB of RAM and 2 bi-GPU NVIDIA Tesla V100 were allocated for the processing task. To give an estimation of the time required to complete the training, the general classifier (CINAC_v1) was trained over 14 epochs. Training took around 40h (less than 3 hours by epoch).

#### Classifier prediction

Using Linux-based workstation with one GPU (NVIDIA® GeForce GTX 1080), 12 CPUs (Intel Xeon CPU W-2135 at 3.70 GHz), and 64 GB of RAM, the time to predict the cell activity on a movie of 12500 frames was on average 13 sec, approximately 3.5 hours for a 1000 cells. The time to predict the cell type on a movie of 12500 frames was on average 2 sec, approximately 33 min for a 1000 cells. Similar performance was achieved using google colab.

### 2.6 Performance evaluation

#### Descriptive metrics for activity classifier: sensitivity, precision, F1 score

We evaluated the performance of the activity classifiers which predict for each frame if a cell is active or not. We chose to measure the sensitivity and precision values, as well as the F1 score that combines precision and sensitivity into a single metric defined as the harmonic mean of precision and sensitivity (Géron, 2019). Because we have a skewed dataset (cells being mostly inactive), we chose not to use the accuracy. In order to compute the metrics, we needed to evaluate the number of true negative and false negative transients. To do this evaluation, we defined all putative transients. We identified the putative transient using the change of the sign of the first derivative on a smooth fluorescence time-course to detect all onsets and peaks. The smoothing is based on the convolution of a scaled Hamming window with the signal using numpy package methods. We considered as a putative transient all segments between an onset and the immediately following peak. Since the output of the binary classifier is the probability for a cell to be active at a given frame, we considered that a transient was predicted as true if at least during one of its frames the cell was predicted as active. On this basis, we were then able to compute the sensitivity (defined as the proportion of real transients that were detected) and the precision (defined as the proportion of detected transients that are real transients).

#### Descriptive metrics for cell type classifier: sensitivity, precision, F1 score

We evaluated the performance of the cell type classifier which predicts the type of a cell. We chose to measure the sensitivity and precision values, as well as the F1 score. To do so we used the metrics module of the Python package scikit-learn (Pedregosa et al., 2011) that returns the confusion matrix and a classification report containing those metrics.

#### Statistical analysis

The distribution of F1 score values on the datasets for each inference method were compared using Wilcoxon signed-rank test with an *a priori* significance level of p = 0.05 using scipy Python package (Oliphant and Jones, 2001). This test was performed only on distribution with more than 15 samples. Significance level: we used ‘*’ for 0.01≤p-value<0.05, ‘**’ for 0.001≤p-value< 0.01 and ‘***’ for p-value<0.001.

#### Detection of overlap activity

For all pairs of overlapping cells (with an intersected area of at least 15% of the highest area of the two cells), we computed their transient profiles over all putative activations (all rise time over the full recording) and then calculated the Pearson correlation with their respective cell source profile. If the correlation was superior to 0.7 for the first cell while inferior to 0.2 in the second one, we considered that the transient was a true activation of the first cell leading to a false transient in the second one. Finally, we evaluated whether the classifier could classify the putative transient of the second cell as false (with a prediction < 0.5).

#### Comparison with CaImAn

We compared the classifier performance against a state of the art computational tool, namely CaImAn. To fairly compare CaImAn and DeepCINAC to the ‘Ground Truth’ we used the cell contours obtained from the CNMF. The spike inference from the MCMC as well as, DeepCINAC predictions and the ‘Ground Truth’ were established on these contours. A transient was considered as detected by CaImAn, if at least one spike was inferred during the rise time of the transient.

### 2.7 DeepCINAC workflow

To summarize, DeepCINAC uses .cinac files built using the GUI. To train a classifier, those files are given as inputs to the neuronal network, providing time series data representing the calcium fluorescence dynamics of the cells. The same files can be used to benchmark the performance of a classifier and using the GUI, it is possible to add new data for training based on the errors of previous classifier outputs (Figure 5).

**Figure 5.**
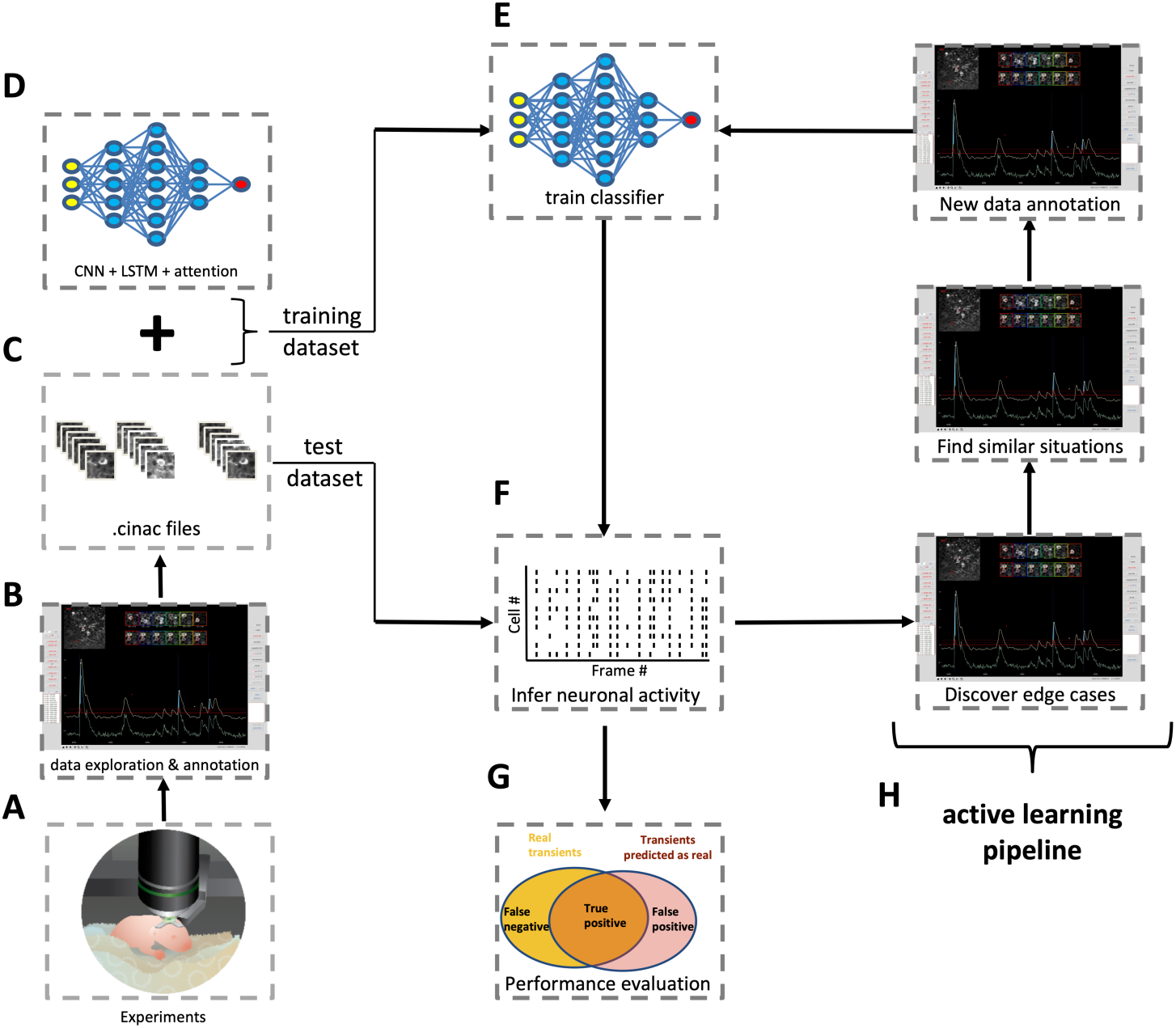
DeepCINAC step by step workflow. **5A**. Schematic of two-photon imaging experiment. **5B**. Screenshot of DeepCINAC GUI used to explore and annotate data. **5C**. The GUI produces .cinac files that contain the necessary data to train or benchmark a classifier. **5D**. Schematic representation of the architecture of the model that will be used to train the classifier and predict neuronal activity. **5E**. Training of the classifier using the previously defined model. **5F**. Schematic of a raster plot resulting from the inference of the neuronal activity using the trained classifier. **5G**. Evaluation of the classifier performance using precision, sensitivity and F1 score. **5H**. Active learning pipeline: screenshots of the GUI used to identify edge cases where the classifier wrongly infer the neuronal activity and annotate new data on similar situations in order to add data for a new classifier training.

### 2.8 Toolbox and data availability

The source code is available on gitlab (https://gitlab.com/cossartlab/deepcinac). The page includes a full description of the method, a user manual, tutorials and test data, as well as the settings used. A notebook configured to work on google colab is also provided, allowing for the classifier to run online, thus avoiding installing the necessary environment and providing a free GPU. The toolbox has been tested on windows (v7 Pro), Mac Os X (MacOS Mojave) and Linux Ubuntu (v.18.04.1).

## 3. Results

### 3.1. Validation of visual Ground truth

As a first step, we asked whether the visualization of fluorescent transients was a good estimation of spiking activity present in a neuron. To do so, we used previously published data combining loose seal cell attached recordings with 2-photon calcium imaging (Chen et al., 2013; GENIE Project, 2015). We compared the visual ‘ground truth’ to the ‘true’ spiking of the cell. We found that visual inspection of calcium imaging movies allows the detection of 87.1, 79.1 and 80.7% ‘true’ transients (i.e. spike associated transient) for each human expert respectively (median sensitivity, Figure 6A). Among visually detected transients, 98.7, 98.6 and 98.6% were ‘true’ transients for each human expert respectively (median precision, Figure 6B). The F1 scores that combine these two previous metrics were 84.1, 81.5, 85.9% for each human expert respectively (median value Figure 6C). We evaluated the classifier CINAC_v6 trained with some recordings of ‘Visual-ctx-6s’ and ‘Hippo-dvt’ (Table S1). We found that it allows the detection of 94% of the ‘true’ transients (median sensitivity, Figure 6A). Among predicted transients, 94.2% were ‘true’ transients (median precision, Figure 6B). F1 score was 94.7% (median value Figure 6C). Overall we conclude that in absence of patch-clamp based ground truth the visual inspection of the movie provides a good estimation of neuronal activity and that deep learning approach based on movie visualization can reach the human level in estimating cell activations.

**Figure 6:**
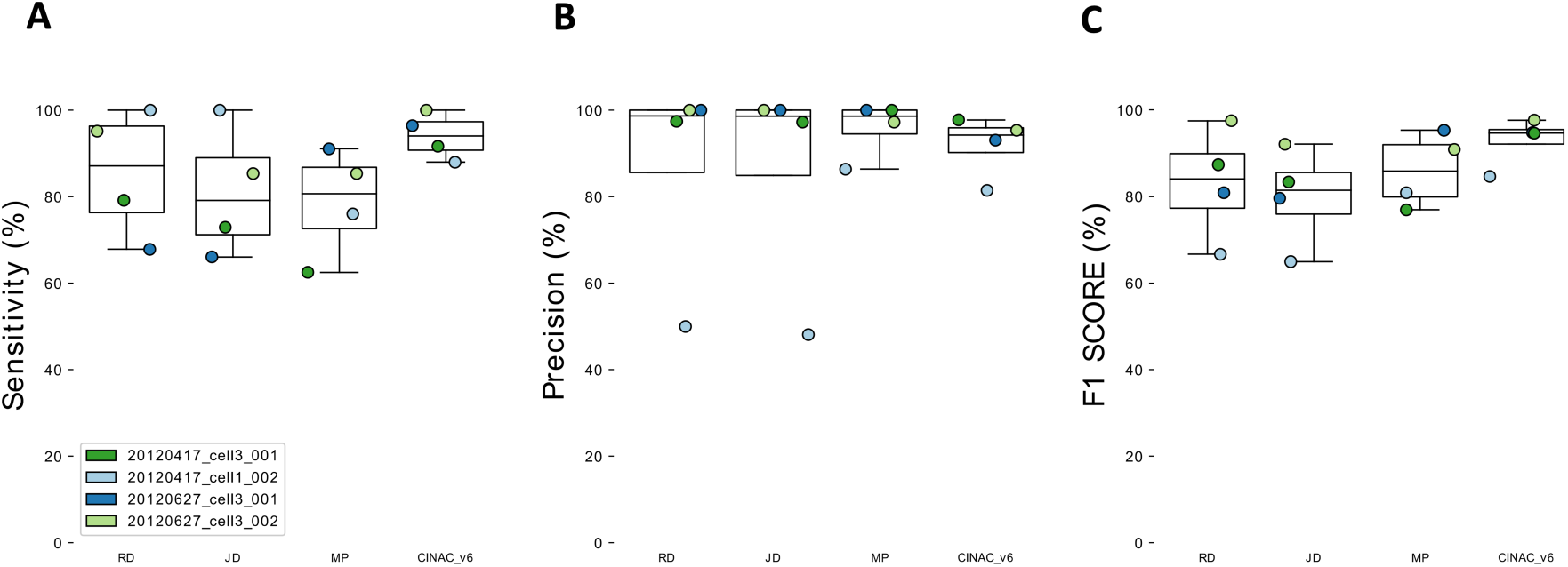
Validation of visual ‘Ground truth’ and deep learning approach. **6A**. Boxplots showing sensitivity for the 3 human experts (RD, JD, MP) and CINAC_v6 evaluated against the known ground truth from 4 cells from the GENIE project. **6B**. Boxplots showing precision for the 3 human experts (RD, JD, MP) and CINAC_v6 evaluated against the known ground truth from 4 cells from the GENIE project. **6C**. Boxplots showing F1 Score for the 3 human experts (RD, JD, MP) and CINAC_v6 evaluated against the known ground truth from 4 cells from the GENIE project. Each colored dot represents a cell. Cell labels in the legend correspond to session identifiers from the dataset. CINAC_v6 is a classifier trained on data from the GENIE project and the ‘Hippo-dvt’ dataset (Table 1, Table S1).

### 3.2 DeepCINAC performance evaluation on developing hippocampus dataset

#### Comparing DeepCINAC against CaImAn and human level

We compared the performance of DeepCINAC and CaImAn (Pnevmatikakis et al., 2016), a well established algorithm to infer neuronal activity, against the visual ground truth on CA1 hippocampus data during development (‘Hippo-dvt’). We first evaluated DeepCINAC (CINAC_v1) on 20 putative pyramidal neurons and 5 interneurons (Figure 7). The median sensitivity was 80.3% (interquartile range 75–94.5, Figure 7A), the median precision was 90.8% (interquartile range 81.2-95.5, Figure 7B) and the median F1 score was 86.3% (interquartile range 78.9-91.3, Figure 7C).

**Figure 7:**
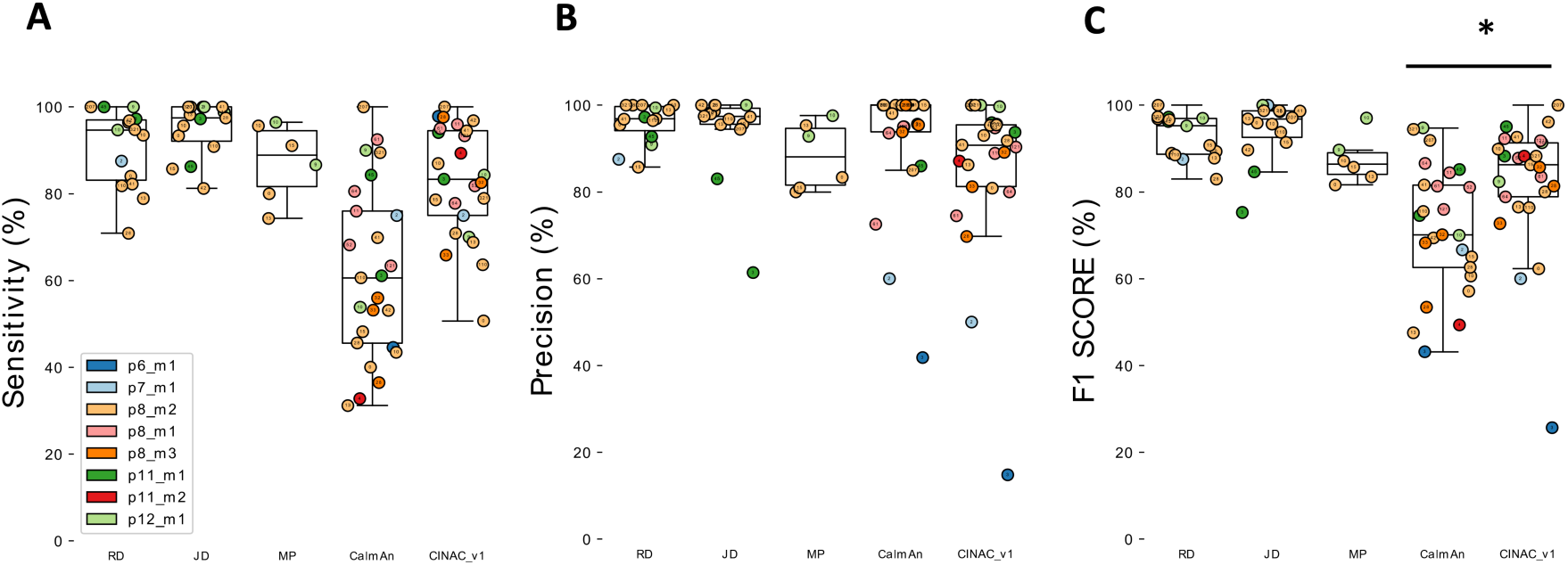
Evaluation of CINAC_v1 performance on ‘Hippo-dvt’ dataset. **7A**. Boxplots showing sensitivity for the 3 human experts (RD, JD, MP), CaImAn and CINAC_v1 evaluated against the visual ground truth of 25 cells. 15 cells were annotated by JD and RD, 6 by MP. **7B**. Boxplots showing precision for the 3 human experts (RD, JD, MP), CaImAn and CINAC_v1 evaluated against the visual ground truth of 25 cells. 15 cells were annotated by JD and RD, 6 by MP. **7C**. Boxplots showing F1 score for the 3 human experts (RD, JD, MP),CaImAn and CINAC_v1 evaluated against the visual ground truth of 25 cells. 15 cells were annotated by JD and RD, 6 by MP. Each colored dot represents a cell, the number inside indicates the cell’s id and each color represents a session as identified in the legend. CINAC_v1 is a classifier trained on data from the ‘Hippo-dvt’ dataset (Table 1, Table S1).

We next evaluated CaImAn on the same cells using the same metrics. The median sensitivity was 60.6% (interquartile range 45.6-76, Figure 7A), the median precision was 100% (interquartile range 93.8-100, Figure 7B). The median F1 score was 70.1% (interquartile range 62.6-81.6, Figure 7C), which was significantly lower than CINAC_v1 F1 score (Wilcoxson signed rank test, T=50 and p=0.002). Finally, we asked if DeepCINAC could perform as well as human ‘experts’. The median CINAC_v1 F1 score on the 15 cells annotated by the 2 human ‘experts’ (JD and RD) was 88.2% (interquartile range 78.3-92) which was significantly lower than RD and JD F1 scores (F1=95.2%, T=4, p=0.002 and F1=96.8%, T=22, p=0.031 respectively, Figure 7-1A). However, on six cells annotated by MP, CINAC_v1 and MP F1 scores were close (F1=84.3% and F1=86.4% respectively, Figure 7-1B). Even though DeepCINAC is still not at the ground truth level (combination of triple human labelling), it approximates human level.

**Figure 7-1:**
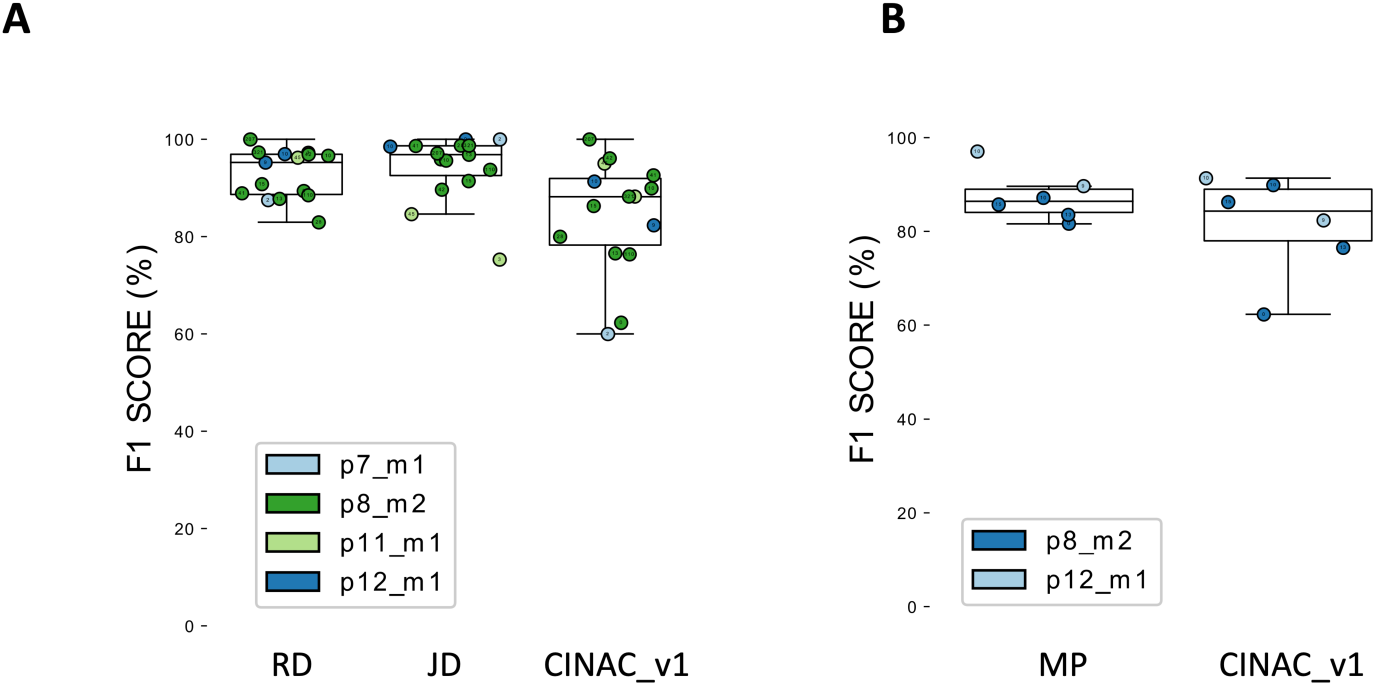
Comparison of CINAC performance to human experts. 7-1A: Boxplot displaying F1 score of two human experts (RD and JD) and CINAC_v1. Here are shown 15 cells annotate by both ‘experts’ 7-1B: Boxplot displaying F1 score of one human expert (MP) and CINAC_v1. Here are shown 6 cells annotated by MP. Each colored dot represents a cell, the number inside indicates the cell’s id and each color represents a session as identified in the legend. CINAC_v1 is a classifier trained on data from the ‘Hippo-dvt’ dataset (Table 1, Table S1).

#### Specific handling of overlap

One important characteristic of data from the developing CA1 region of the hippocampus is the high density of active neurons that can lead to overlap. This overlap between cells leading to false transients was pointed out as a specific issue in the analysis of calcium traces from a demixing (Gauthier et al., 2018). We asked whether the classifier would be able to distinguish real transients from increases in fluorescence due to the activity of an overlapping cell. Based on the visual inspection of imaged fields of view with numerous overlaps, we chose to specifically test the algorithm on calcium imaging data containing 391 cells segmented using CaImAn. Among those cells, we detected a total of 426 transients (fluorescence rise time) from 23 cells that were likely due to overlapping activity from a neighboring cell (see method for overlap activity detection). Among those transients, 98.6% were correctly classified as false by CINAC_v1 (general classifier), 93.2% were correctly classified as false by the CINAC_v7 (interneuron specific classifier) and 93.2% were correctly classified as false by CaImAn. We next asked if the results could be improved by the use of another segmentation method. To do so, we performed the same analysis on the exact same field of view using the classifier prediction on the segmented cells obtained from suite2p (Pachitariu et al., 2017). Among a total of 480 cells, a total of 2718 transients from 101 cells were likely due to the activation of an overlapping cell, 99.1% of them were correctly classified as false by CINAC_v1.

#### Onset to peak prediction

Since we aimed at predicting as active all the frames included in the full rise time of the calcium transient (from onset to peak), we looked at the proportion of frames predicted as active in real transients. Using the general classifier (CINAC_v1), the median ratio of frames predicted among each real transient was 85.7% (interquartile range 70-100) for the 20 putative pyramidal cells and the 5 putative interneurons (Figure 7-2). We demonstrated that CINAC_v1 allows the detection of cell activation all along the rise time, giving us both the onset of cell activation and the duration of the rise time (Figure 7-2).

**Figure 7-2:**
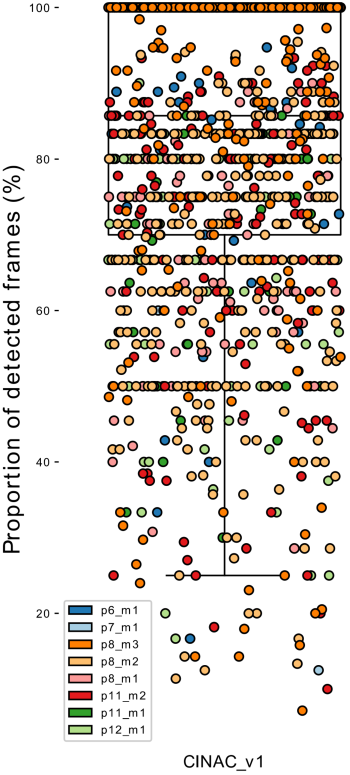
Onset to peak detection of calcium transient. Boxplot showing the proportion of frames predicted as active during the transient rise time. CINAC_v1 is a classifier trained on data from the ‘Hippo-dvt’ dataset (Table 1, Table S1). Each colored dot represents a transient and each color represents a session as identified in the legend.

### 3.3 Classifier generalization and specialization

#### DeepCINAC performances on other datasets

A major aspect to consider in the development of algorithms to infer neuronal activity from calcium imaging data, is the ability to be easily scalable to the wide variety of datasets (i.e. different indicators, different brain regions,…).

We investigated the extent to which DeepCINAC (CINAC_v1), that was trained on data from the developing hippocampus, would perform on other datasets (Figure 8-1). To answer that question, we used i) GECO imaging movies (‘Hippo-GECO’, 5 Hz, 2 µm/pixel, Figure 8-1A), ii) GCaMP6m imaging movies (‘Hippo-6m’, 10 Hz, 2 µm/pixel, Figure 8-1B) both from the adult hippocampus, iii) GCAMP6s calcium imaging movies from the developing barrel cortex (‘Barrel-ctx-6s’, 1.5Hz, 1.2µm/pixel, Figure8-1C) (Modol et al., 2019) iv) GCAMP6s calcium imaging movies of interneurons from the developing hippocampus (‘Hippo-dvt-INs’, 10 Hz, 2 µm/pixel,, Figure 8-1D) and GCaMP6s recordings from the adult visual cortex (‘Visual-ctx-6s’ - downsampled 10Hz, rescaled 1.2 µm/pixel, see methods, Figure 8-2). We show that DeepCINAC performs well on ‘Hippo-6m’ and ‘Barrel-ctx-6s’ data. On ‘Hippo-6m’, F1 scores were 66.7% and 70.9% for CaImAn and CINAC_v1 respectively (Figure 8-1B, bottom panel). On ‘Barrel-ctx-6s’, F1 scores were 54.3% and 76.4% for CaImAn and CINAC_v1 respectively (Figure 8-1C, bottom panel). However CINAC_v1 does not generalize well enough to infer activity on the ‘Hippo-GECO’ recordings (F1 score = 44.2%, Figure 8-1A) neither on ‘Visual-ctx-6s’ (F1 score = 69.9%, Figure 8-1C).

**Figure 8:**
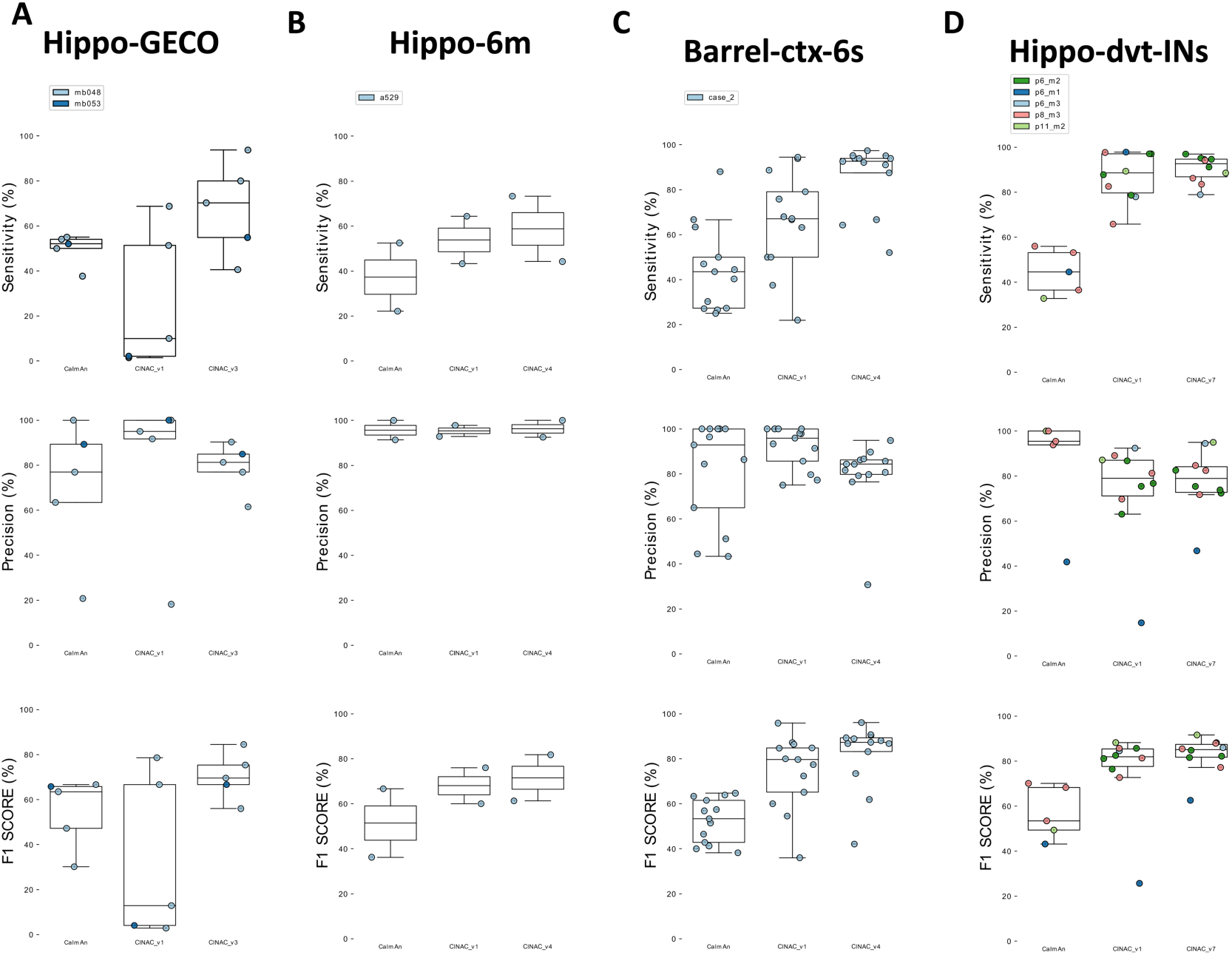
Use of DeepCINAC classifiers to optimize performances on various dataset. **8A**: Boxplot displaying the sensitivity (top panel), precision (middle panel) and F1 score (bottom panel) for ‘Hippo-GECO’ dataset. For each panel, we evaluated CaImAn performance as well as two different versions of CINAC (v1 and v3). CINAC_v1 is a classifier trained on data from the ‘Hippo-dvt’ dataset and CINAC_v3 is a classifier trained on data from the ‘Hippo-GECO’ dataset (Table 1, Table S1). **8B**: Boxplot displaying the sensitivity (top panel), precision (middle panel) and F1 score (bottom panel) for ‘Hippo-6m’ dataset. For each panel, we evaluated CaImAn performance as well as two different versions of CINAC (v1 and v4). CINAC_v1 is a classifier trained on data from the ‘Hippo-dvt’ dataset and CINAC_v4 is a classifier trained on data from the ‘Hippo-dvt’, ‘Hippo-6m’, and ‘Barrel-ctx-6s’ dataset (Table 1, Table S1). **8C**: Boxplot displaying the sensitivity (top panel), precision (middle panel) and F1 score (bottom panel) for ‘Barrel-ctx-6s’ dataset. For each panel, we evaluated CaImAn performance as well as two different versions of CINAC (v1 and v4). CINAC_v1 is a classifier trained on data from the ‘Hippo-dvt’ dataset and CINAC_v4 is a classifier trained on data from the ‘Hippo-dvt’, ‘Hippo-6m’, and ‘Barrel-ctx-6s’ dataset (Table 1, Table S1). **8D**: Boxplot displaying the sensitivity (top panel), precision (middle panel) and F1 score (bottom panel) for ‘Hippo-dvt-INs’ dataset. For each panel, we evaluated CaImAn performance as well as two different versions of CINAC (v1 and v7). CINAC_v1 is a classifier trained on data from the hippo-dvt dataset and CINAC_v7 is a classifier trained on interneurons from the ‘Hippo-dvt’ dataset (Table 1, Table S1). Each colored dot represents a cell, the number inside indicates the cell’s id and each color represents a session as identified in the legend.

**Figure 8-1:**
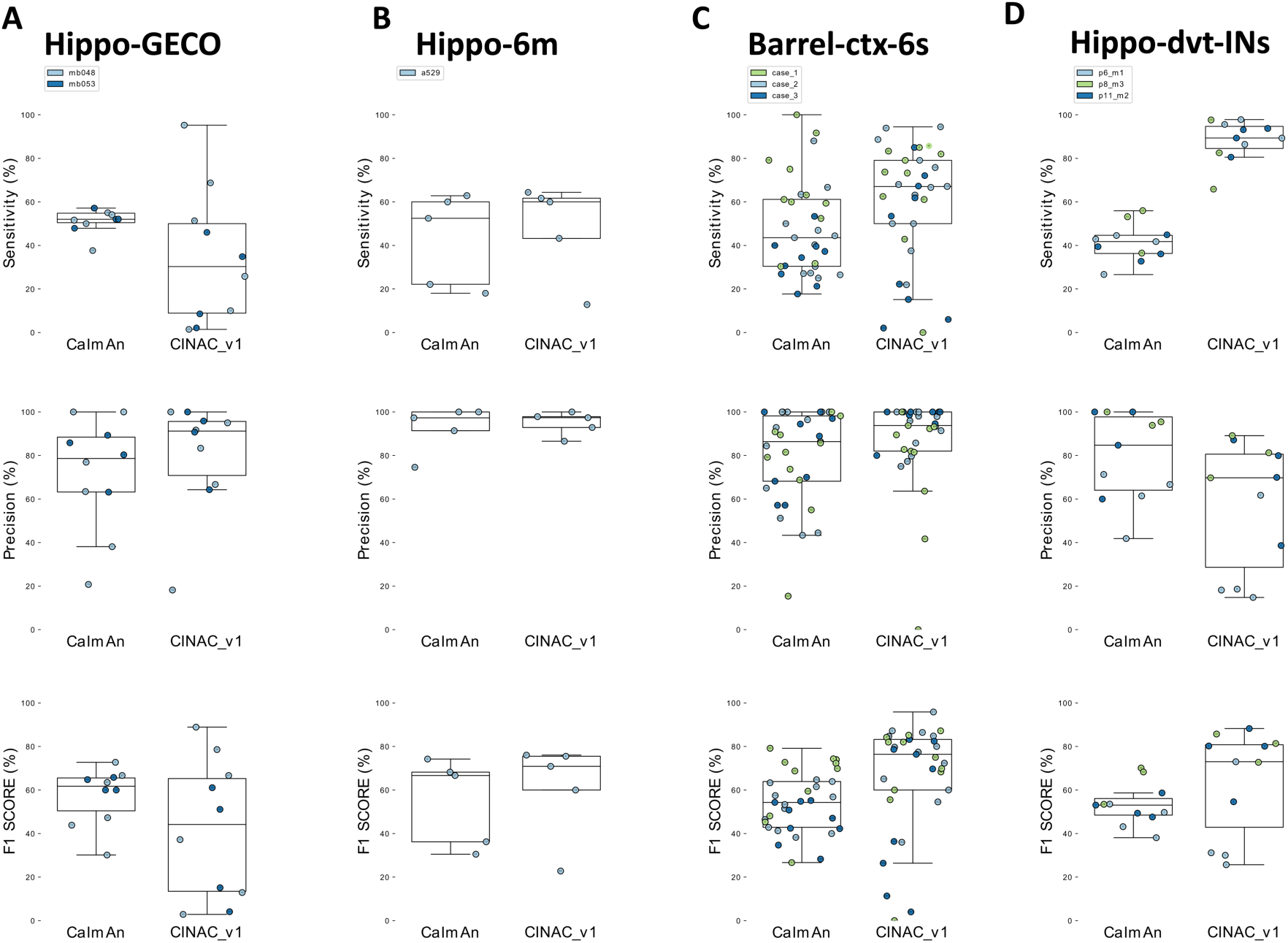
Comparison of CaImAn and CINAC_v1 performances on various dataset. **8-1A:** Boxplot displaying the sensitivity (top panel), precision (middle panel) and F1 score (bottom panel) for ‘Hippo-GECO’ dataset. For each panel, we evaluated CaImAn performance as well as CINAC_v1. **8-1B:** Boxplot displaying the sensitivity (top panel), precision (middle panel) and F1 score (bottom panel) for ‘Hippo-6m’ dataset. **8-1C:** Boxplot displaying the sensitivity (top panel), precision (middle panel) and F1 score (bottom panel) for ‘Barrel-ctx-6s’ dataset. **D:** Boxplot displaying the sensitivity (top panel), precision (middle panel) and F1 score (bottom panel) for ‘Hippo-dvt-INs’ dataset. Each colored dot represents a cell, the number inside indicates the cell’s id and each color represents a session as identified in the legend. CINAC_v1 is a classifier trained on data from the ‘Hippo-dvt’ dataset (Table 1, Table S1).

**Figure 8-2:**
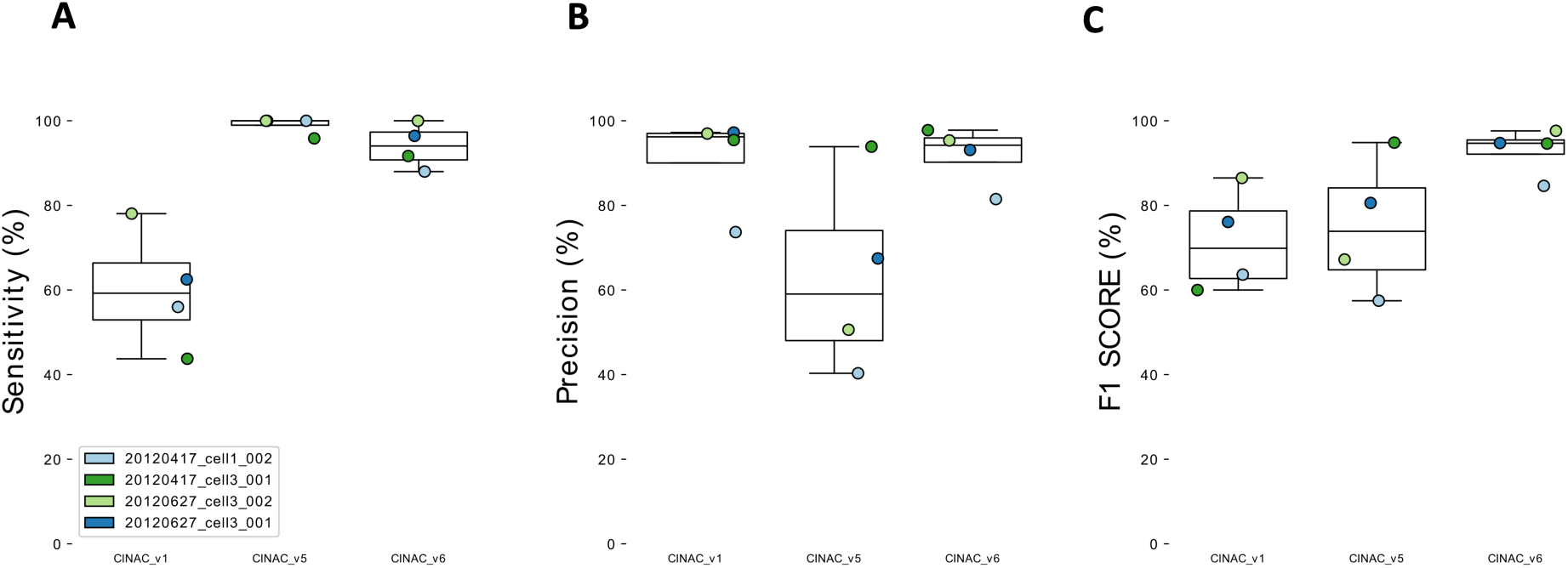
Use of DeepCINAC classifiers to optimize performances on ‘Visual-ctx-6s’ dataset. **A**. Boxplots showing sensitivity for CINAC_v1, CINAC_v5 and CINAC_v6 evaluated against the known ground truth of 4 cells from the GENIE project. **8-2B**. Boxplots showing precision for CINAC_v1, CINAC_v5 and CINAC_v6 evaluated against the known ground truth of 4 cells from the GENIE project. **8-2C**. Boxplots showing F1 Score for CINAC_v1, CINAC_v5 and CINAC_v6 evaluated against the known ground truth of 4 cells from the GENIE project. CINAV_v1 is a classifier trained on data from the ‘Hippo-dvt’ dataset, CINAC_v5 is a classifier trained on data from ‘Visual-ctx-6s’ dataset, CINAC_v6 is a classifier trained on data from ‘Visual-ctx-6s’ dataset and 4 cells from the ‘Hippo-dvt’ dataset (Table 1, Table S1). Each colored dot represents a cell. Cell labels in the legend correspond to session identifiers from the dataset.

To overcome these poor performances on ‘Hippo-GECO’ and ‘Visual-ctx-6s’ and to improve performances on ‘Barrel-ctx-6s’ and ‘‘Hippo-6m’ datasets, we considered two strategies. The first one consists in training a classifier specific to the data. The second one consists in adding part of the new data to the large database to improve the classifier ability to generalize (Figure 8 and 8-1).

Since our performances were low using CINAC_v1, we adopted the first strategy to improve the classifier on ‘Hippo-GECO’ and ‘Visual-ctx-6s’ datasets. We used part of these datasets to train specific classifiers and evaluate them on the remaining data (Table S1). First, we observed that the ‘Hippo-GECO’ specific classifier (i.e. CINAC_v3) performed better than CINAC_v1 and CaImAn (CINAC_v3 median F1 score=69.6%, CINAC_ v1 median F1 score=12.9% and CaImAn median F1 score=63.5%, Figure 8A bottom panel). This increase in F1 score from CINAC_v1 to CINAC_v3 was due to an increase in the sensitivity of the classifier (CINAC_v1 median sensitivity=10%, CINAC_v3 median sensitivity=70.3%, Figure 8A top panel) with a moderate loss in precision (CINAC_v1 median precision=95%, CINAC_v3 median precision= 81.2%, Figure 8A middle panel). Second, we showed that the ‘Visual-ctx-6s’ datasets specific classifiers (i.e. CINAC_V5 and CINAC_v6) performed better than CINAC_v1 and CaImAn (CINAC_v1 median F1 score=69.9%, CINAC_v5 median F1 score=73.9% and CINAC_v6 median F1 score=94.7%, Figure 8-2C). Because CINAC_v5 was trained exclusively on labelled data from the ‘visual-ctx-6s’ dataset it allows an increase in the classifier sensitivity (CINAC_v1 median sensitivity=59.2%, CINAC_v5 median sensitivity=100%, Figure 8-2A). However this increase was achieved at the cost of a reduced precision (CINAC_v1 median precision=96.2%, CINAC_v5 median precision=59%, Figure 8-2B), and overall a slight increase in the F1 score (Figure 8-2 C). To improve the performance on this dataset, we extended the CINAC_v5 training set with 4 cells from the ‘Hippo-dvt’ to train CINAC_v6. This allows us to increase both sensitivity (CINAC_v6 median sensitivity=94%) and precision (CINAC_v6 median precision=94.2%) of the classifier, leading to a large improvement of the F1 score (CINAC_v6 median F1 score=94.7%, Figure 8-2C). In a nutshell, when the dataset to analyse has different calcium dynamics, a new classifier specifically trained on this dataset would reach higher performance than CINAC_v1.

On the datasets from the adult hippocampus (‘Hippo-6m’) and developing barrel cortex (‘Barrel-ctx-6s’), since CINAC_v1 performances were close to the performance reached on ‘Hippo-dvt’ dataset, we considered the second strategy. We extended the CINAC_v1 training dataset with labeled data from ‘Hippo-6m’ and ‘Barrel-ctx-6s’ datasets and trained a new classifier (CINAC_v4). First, we observed that on the ‘Hippo-6m’ dataset, CINAC_v4 classifier has better sensitivity and precision than CINAC_v1 and CaImAN (CINAC_v1 median sensitivity=53.8%, CINAC_v4 median sensitivity=58.7%, CaImAn median sensitivity=37.3%, CINAC_v1 median precision=95.3%, CINAC_v4 median precision=96.2%, CaImAn median precision=95.7%, CINAC_v1 median F1 score=68%, CINAC_v4 median F1 score=71.6%, CaImAn median F1 score=51.4%, Figure 8B). Second, we found that on ‘Barrel-ctx-6s’ dataset, CINAC_v4 classifier performed better than CINAC_v1 and CaImAn (CINAC_v4 F1 score=87.3%, CINAC_v1 F1 score=79.7%, CaImAn F1 score=53.3%, Figure 8C, bottom panel). This increase in F1 score from CINAC_v1 to CINAC_V4 was due to an increase in the sensitivity of the classifier (CINAC_v4 median sensitivity=92.6%, CINAC_v1 median sensitivity=67.1%, CaImAn median sensitivity=43.5%, Figure 8C, top panel) with a moderate loss in precision (CINAC_v4 median precision=84.3%, CINAC_v1 median precision=95.9%, CaImAn median precision=92.9%, Figure 8C, middle panel). Overall we confirm here that adding part of a new dataset to the training set of CINAC_v1 classifier allows us to improve the performance on this new dataset. Thus, one could use the training dataset of CINAC_v1 and add part of a new dataset to train a classifier that would achieve better performance than CINAC_v1 on this new data.

Because we benefited from already published data from our group (‘Barrel-ctx-6s’), we next asked if we could arrive at the same conclusion using CINAC_v1. We used the activity inferred by CINAC_v1 on this data and performed assemblies detection analysis as described previously (Modol et al., 2019). We found the same number of assemblies as the original analysis (Figure 8-3A and 8-3B, first two panels), as well as the same topographic organization (Figure 8-3A and 8-3B, bottom panels). We confirmed that the assemblies detected by either CaImAn or CINAC_v1 were composed of similar cells (Figure 8-3C).

**Figure 8-3:**
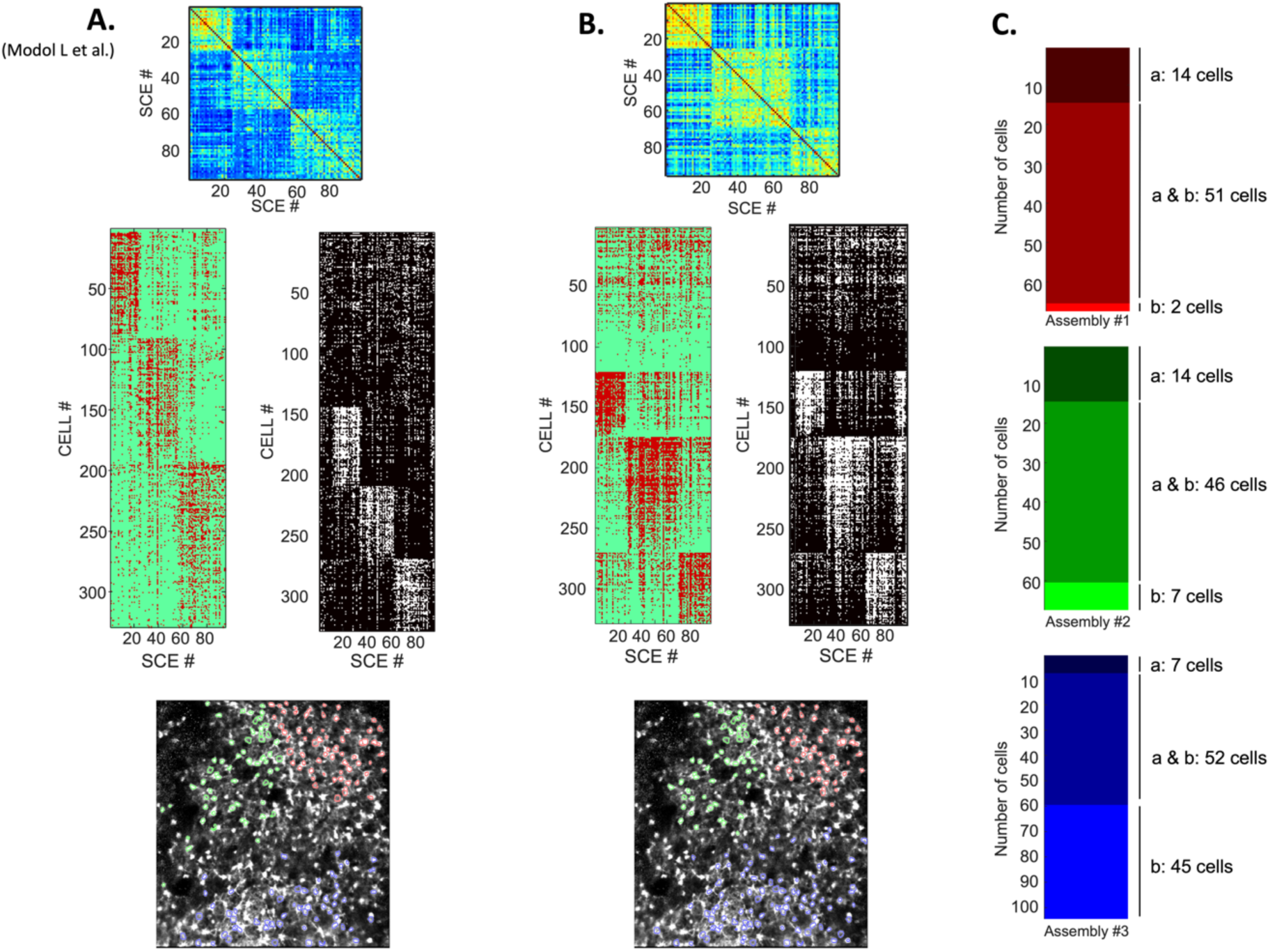
Cell assemblies detection and organization using CaImAn and CINAC_v1 on published data. **8-3A&B**. The top panel represents the clustered covariance matrix of synchronous calcium events (SCE). The middle panel represents neurons active in SCE organized by cluster (cell assembly). The bottom panel represents the cell’s map, each color represents a cell assembly. **8-3A**. Cell assemblies detection results using CaImAn. **8-3B**. Cell assemblies detection results using CINAC_v1. **8-3C**. Individual cells composing assemblies in each method. ‘a’ represents the number of neurons detected by Modol et al., using CaImAn. ‘b’ represents the number of neurons detected using CINAC_v1. Each color represents a cell assembly, color-coded as in the maps.

#### DeepCINAC performances on different cell types

A second important aspect to infer neuronal activity from calcium imaging movies is the variety of cell types recorded in the same field of view (e.g. interneuron and pyramidal cells). In recordings from the hippocampus we observed that most interneurons have very different calcium dynamics than pyramidal cells (higher fluorescence signal followed by a plateau). Because CINAC_v1 was mainly trained on the activity of pyramidal cells, we suspected that it would not provide accurate inference on interneurons. Using the GUI we verified its prediction on interneurons and concluded that they were not always optimal (Figure 8-1D). To improve activity inference on interneurons, we trained an interneuron specific classifier. In more details the precision of the inference was similar for CINAC_v1 and CINAC_v7 (CINAC_v1 median precision=79%, CINAC_v7 median precision=78.9%, Figure 8D top panel). However CINAC_v7 provides more sensitive inference (CINAC_v1 median sensitivity=88.6%, CINAC_v7 median sensitivity=92.6%, Figure 8D middle panel). As a result, the specific classifier performed better than the general one on interneurons (CINAC_v1 median F1 score=81.9%, CINAC_v7 median F1 score=85.1%, Figure 8D, Bottom panel).

### 3.4 Cell type inference using DeepCINAC

Recently a deep-learning method using a similar model to DeepCINAC was proposed to differentiate cell types (Troullinou et al., 2019). This model was based on the analysis of fluorescence traces from various cell types and automatically classified imaged cells in different types. We asked whether DeepCINAC would be able to distinguish interneurons from pyramidal cells using as input the calcium imaging movie rather than the fluorescence trace. Additionally we added a noise category in the training dataset allowing us to automatically discard cells. We achieved a general F1 score of 86%. We had a sensitivity of 90.2%, 81.6% and 81.8% and a precision of 90.2%, 91.2% and 60% for pyramidal, interneuron and noisy cells respectively (Table 2).

**Table 2:** Cell type prediction confusion matrix. Confusion matrix, representing the number of True Positives, True Negatives, False Positives and False negatives. “Ground Truth” refers to the manually detected interneurons and pyramidal cells. “Prediction” refers to the type predicted by the classifier for the same cells.

|  |  | Ground Truth |  |  |
| --- | --- | --- | --- | --- |
|  |  | Pyramidal cell | Interneuron | Noise |
| Prediction | Pyramidal cell | **46** | 5 | 0 |
|  | Interneuron | 1 | **31** | 2 |
|  | Noise | 4 | 2 | **9** |

Since activity inference performance using DeepCINAC depends on the cell type, we perform this cell type prediction before activity inference. During the activity inference of a movie DeepCINAC can be configured to switch between different activity classifiers depending on the type of the cell to predict.

## 4. Discussion

Deep learning based method(s) to infer neuronal activity from 2-photon calcium imaging datasets use cellular fluorescence signals as inputs. Here we propose a method based on the visual inspection of the recordings. We will discuss the advantages and limitations of this approach.

Using the movie dynamics, we benefited from all the information available in the calcium imaging movie. This approach allowed us to not rely on a demixing algorithm to produce the neuron’s traces. Instead, by working directly on the raw calcium imaging movie, the algorithm has learned to identify a transient and distinguish overlap activity from a real transient. DeepCINAC achieves better performance than CaImAn and is able to achieve human performance level on some fields of view and cells.

Additionally, we show that a classifier trained on a specific dataset (‘Hippo-dvlt-6s’) is able to generalize to other datasets (‘Hippo-6m’ and ‘Barrel-ctx-6s’). DeepCINAC allows training of flexible classifiers whose generalization on new datasets can be improved by adding part of this new dataset to the training (at the cost of slightly reduced performance of the classifier on original data). However, we show that generalization is not always achieved such as in the case of a classifier trained on ‘Hippo-dvt’ data and used to predict activity on some very different datasets (‘Hippo-GECO’ and ‘Visual-ctx-6s’). This is likely explained by the difference in calcium indicator, imaging rate and imaging resolution. We demonstrated that this limitation can be circumvented by training specific classifiers. Overall, this approach allowed us to create classifiers that scale to different developmental stages (from P5 to adult), different types of neurons (pyramidal cells and interneurons) as well as different indicators (GCaMP6s, GCaMP6m, GECO).

On a side note, we observed during the visual inspection of the prediction through the GUI that the network might be able to handle fake transients due to X and Y movements or neuropil activation. By avoiding the use of any threshold to select transients over fluorescence time-course, the classifier might also be able to detect: i) small amplitude transients, ii) transients occurring during the decay of another one, iii) summations. Importantly, because activity is split into segments of 100 frames (around 10 sec), DeepCINAC classifiers would still perform well and deal with changes in fluorescence baseline.

Finally, we explored the range of values of hyperparameters in order to optimize the accuracy of the classifier. Labeling data is time-consuming but the training does not need any parameters tuning and the prediction is straight forward. Neither tedious manual tuning of parameters is required, nor a GPU on a local device because we provide a notebook to run predictions on google colab (see Methods). Predictions are fast, with a run-time of around 10 seconds by cell for 12500 frames, meaning less than three hours hours for 1000 cells. However, a GPU would be necessary to train the network on a big dataset.

Already widely used by many calcium imaging labs (Andalman et al., 2019; Driscoll et al., 2017; Gauthier and Tank, 2018; Katlowitz et al., 2018), CaImAn offers a performing and functional analysis pipeline. Even though the complex fine tuning of CaImAn parameters on the dataset might lead to a suboptimal spike inference from the model, we decided to compare CaImAn against our ground truth.

The benchmarks remain limited to a small number of cells for which we established a ground truth and may be extended to more cells. Notably, a future approach could be to use more realistic simulated data such as done in a recent work (Charles et al., 2019).

In the model we used, each cell was represented by a segment of the field of view, in our case a 25 by 25 pixels (50 µm by 50 µm) window that allows complete coverage of the cell fluorescence and potential overlapping cells. Consequently, the network is able to generalize its prediction to recordings acquired with this resolution (2 µm / pixel). However, to be efficient on another calcium imaging dataset with a different resolution it would be necessary to train a new classifier adjusting the window size accordingly. Importantly, we trained the model on a selection of cells with valid segmentations; meaning that a cell is not represented by several contours. The inference performance of the classifier might decrease on cells whose segmentation was not properly achieved.

Since precise spike inference cannot be experimentally assessed on the data, we chose to infer the activity of the cell defined by the fluorescence rise time instead of inferring the spikes. However, with a ground truth based on patch-clamp recordings, we could adapt this method to switch from a binary classification task to a regression task, predicting the number of spikes at each frame.

## Conclusion

We built DeepCINAC basing the ground truth on visual inspection of the movie and training the classifier on movie segments. DeepCINAC offers a flexible, fast and easy-to-use toolbox to infer neuronal activity from a variety of two photon calcium imaging dataset, reaching human level performance. It provides the tools to measure its performance based on human evaluation. Currently, DeepCINAC provides several trained classifiers on CA1 two-photon calcium imaging at early postnatal stages; its performance might still be improved with more labeled data. In the future, we believe that a variety of classifiers collaboratively trained for specific datasets should be available to open access.

## Video Legends

***Video 1:*** *In vivo 2-p imaging in the CA1 region of the hippocampus in a 12 days old mouse pup*.

*FOV is 80 µm by 80 µm, frame rate is 8Hz and video is speeded up 10 times. The Video shows recurrent periods of neuronal activations recruiting a large number of adjacent neurons leading to spatial and temporal overlaps*.

***Video 2:*** *In vivo 2-p imaging in the CA1 region of the hippocampus in a 7 days old mouse pup*.

*FOV is 100 µm by 100 µm, frame rate is 8Hz and video is speeded up 10 times. The Video shows different cell types (i*.*e. interneurons and pyramidal cells) with different calcium dynamics*.

## Author Contributions

JD, RD, RC and MP Designed Research; RD and JD Performed Research; JD Wrote code; MP JD RD EQ Labeled data. RD, JD, MP, RC Wrote the paper.

## Acknowledgments and Funding source

We would like to thank Marco Bocchio, Laura Modol, Yannick Bollmann, Susanne Reichinnek and Vincent Villette for providing us calcium imaging data.

“Centre de Calcul Intensif d’Aix-Marseille” is acknowledged for granting access to its high performance computing resources. This work was supported by the European Research Council under the European Union’s FP7 and Horizon 2020 research and innovation program (grants no. 242842 and 646925). J.D. was supported by the Fondation pour la Recherche Médicale (grant no. FDM20170638339). M.P was supported by the Fondation pour la Recherche Médicale (grant no. ARF20160936186).

