## Supplementary material for "DeepCINAC: a deep-learning-based Python toolbox for inferring calcium imaging neuronal activity based on movie visualization": Table S1

***Table S1:*** *Detailed data used to train and test the classifiers.*

*Detailed content of training and test datasets used for all CINAC versions (v1 to v7) used in the analysis.*

| CINAC version | Session id | Cell id | n frames | Dataset(s) | training | Testing (figures) |
|---|---|---|---|---|---|---|
| v1 | p5_m1 | 3, 42, 43, 48 | 6400 | Hippo-dvt | yes | no |
| v1 | p7_m1 | 3, 7, 8, 10, 11, 12, 14, 15, 17, 18, 24, 25, 40, 50, 58, 110 | 130523 | Hippo-dvt | yes | no |
| v1 | p7_m2 | 15, 20, 21, 30 | 5158 | Hippo-dvt | yes | no |
| v1 | p7_m3 | 10 | 1600 | Hippo-dvt | yes | no |
| v1 | p8_m2 | 1, 2, 5, 6, 9 | 24791 | Hippo-dvt | yes | no |
| v1 | p8_m3 | 0, 1, 6, 7, 9, 10, 11, 18, 24 | 112500 | Hippo-dvt-INs | yes | no |
| v1 | p10_m1 | 12, 14 | 4200 | Hippo-dvt | yes | no |
| v1 | p12_m1 | 0, 3, 6, 7, 12, 14, 15, 19 | 100000 | Hippo-dvt | yes | no |
| v1 | p13_m1 | 0, 2, 5, 12, 13, 31, 42,44, 48, 51 | 125000 | Hippo-dvt | yes | no |
| v1 | p16_m1 | 9, 55, 62 | 2800 | Hippo-dvt | yes | no |
| v1 | p11_m1 | 6, 11, 12, 17, 22, 24, 25, 29, 30, 33 | 96300 | Hippo-dvt | yes | no |
| v1 | sim_1 | 0, 11, 22, 31, 38, 43, 56, 64, 70, 79, 86, 96, 110, 118, 131, 136 | 40000 | Hippo-dvt | yes | no |
| v1 | sim_2 | 0, 9, 18, 26, 34, 41, 46, | 40000 | Hippo-dvt | yes | no |

| | | | | | | |
|---|---|---|---|---|---|---|
| | | 56, 62, 77, 88, 101, 116, 127, 140, 150 | | | | |
| v1 | p7_m1 | 2, 25 | 25000 | Hippo-dvt | no | **7, 7-1, 7-2** |
| v1 | p8_m1 | 11, 52, 61, 64, 121 | 62500 | Hippo-dvt | no | **7, 7-2** |
| v1 | p8_m2 | 0, 10, 13, 15, 28, 41, 42, 110, 207, 321 | 125000 | Hippo-dvt | no | **7, 7-1, 7-2** |
| v1 | p12_m1 | 9, 10 | 25000 | Hippo-dvt | no | **7, 7-1, 7-2** |
| v1 | p11_m1 | 3, 45 | 25000 | Hippo-dvt | no | **7, 7-1, 7-2** |
| v3 | mb048 | 1, 11 | 15000 | Hippo-GECO | yes | no |
| v3 | mb053 | 14, 16, 30 | 30000 | Hippo-GECO | yes | no |
| v3 | mb048 | 15, 22, 33, 47 | 30000 | Hippo-GECO | no | **8A** |
| v3 | mb053 | 141 | 10000 | Hippo-GECO | no | **8A** |
| v4 | a529 | 0, 31, 64 | 42000 | Hippo-6m | yes | no |
| v4 | case1 | 0, 1, 4, 5, 6, 50, 52, 56, 91, 107, 126 | 19800 | Barrel-ctx-6s | yes | no |
| v4 | case3 | 0, 1, 3, 4, 5, 104, 153, 205, 260 | 16200 | Barrel-ctx-6s | yes | no |
| v4 | p5_m1 | 3, 42, 43, 48 | 6400 | Hippo-dvt | yes | no |
| v4 | p7_m1 | 3, 7, 8, 10, 11, 12, 14, 15, 17, 18, 24, 25, 40, 50, 58, 110 | 130523 | Hippo-dvt | yes | no |
| v4 | p7_m2 | 15, 20, 21, 30 | 5158 | Hippo-dvt | yes | no |
| v4 | p7_m3 | 10 | 1600 | Hippo-dvt | yes | no |
| v4 | p8_m2 | 1, 2, 5, 6, 9 | 24791 | Hippo-dvt | yes | no |
| v4 | p8_m3 | 0, 1, 6, 7, 9, 10, 11, 18, | 112500 | Hippo-dvt, Hippo-dvt-INs | yes | no |

| | | | | | | |
|---|---|---|---|---|---|---|
| | | 24 | | | | |
| v4 | p10_m1 | 12, 14 | 4200 | Hippo-dvt | yes | no |
| v4 | p12_m1 | 0, 3, 6, 7, 12, 14, 15, 19 | 100000 | Hippo-dvt | yes | no |
| v4 | p13_m1 | 0, 2, 5, 12, 13, 31, 42,44, 48, 51 | 125000 | Hippo-dvt | yes | no |
| v4 | p16_m1 | 9, 55, 62 | 2800 | Hippo-dvt | yes | no |
| v4 | p11_m1 | 6, 11, 12, 17, 22, 24, 25, 29, 30, 33 | 96300 | Hippo-dvt | yes | no |
| v4 | sim_1 | 0, 11, 22, 31, 38, 43, 56, 64, 70, 79, 86, 96, 110, 118, 131, 136 | 40000 | Hippo-dvt | yes | no |
| v4 | sim_2 | 0, 9, 18, 26, 34, 41, 46, 56, 62, 77, 88, 101, 116, 127, 140, 150 | 40000 | Hippo-dvt | yes | no |
| v4 | a529 | 12, 27 | 28000 | Hippo-6m | no | **8B** |
| v4 | case2 | 0, 21, 32, 35, 41, 100, 122, 130, 134, 202, 203, 254, 260 | 23400 | Barrel-ctx-6s | no | **8C** |
| v5 | 20120416 | cell1_001, cell1_002 | 3600 | Visual-ctx-6s | yes | no |
| v5 | 20120417 | cell3_002 cell3_003, cell4_001, cell4_002, cell4_003, cell5_002 | 13400 | Visual-ctx-6s | yes | no |
| v5 | 20120515 | cell1_003, cell1_004, | 9600 | Visual-ctx-6s | yes | no |

| | | | | | | |
|---|---|---|---|---|---|---|
| | | cell1_005, cell1_006 | | | | |
| v5 | 20120627 | cell4_002, cell4_004, cell4_005 | 7200 | Visual-ctx-6s | yes | no |
| v5 | 20120417 | cell_1_002, cell3_001 | 4800 | Visual-ctx-6s | no | **8-2** |
| v5 | 20120627 | cell3_002 | 2400 | Visual-ctx-6s | no | **8-2** |
| v5 | 20120627 | cell3_001 | 2400 | Visual-ctx-6s | no | **8-2** |
| v6 | 20120416 | cell1_001, cell1_002 | 3600 | Visual-ctx-6s | yes | no |
| v6 | 20120417 | cell3_002 cell3_003, cell4_001, cell4_002, cell4_003, cell5_002 | 13400 | Visual-ctx-6s | yes | no |
| v6 | 20120515 | cell1_003, cell1_004, cell1_005, cell1_006 | 9600 | Visual-ctx-6s | yes | no |
| v6 | 20120627 | cell4_002, cell4_004, cell4_005 | 7200 | Visual-ctx-6s | yes | no |
| v6 | p12_m1 | 0, 3, 7, 14 | 50000 | Hippo-dvt | yes | no |
| v6 | 20120417 | cell_1_002, cell3_001 | 4800 | Visual-ctx-6s | no | **6, 8-2** |
| v6 | 20120627 | cell3_002 | 2400 | Visual-ctx-6s | no | **6, 8-2** |
| v6 | 20120627 | cell3_001 | 2400 | Visual-ctx-6s | no | **6, 8-2** |
| v7 | p6_m1 | 0, 1, 2 | 37500 | Hippo-dvt-INs | yes | no |
| v7 | p6_m4_a | 0, 11 | 25000 | Hippo-dvt-INs | yes | no |
| v7 | p6_m4_b | 0, 6 | 25000 | Hippo-dvt-INs | yes | no |
| v7 | p6_m5_a | 5, 27 | 25000 | Hippo-dvt-INs | yes | no |
| v7 | p6_m5_b | 2 | 12500 | Hippo-dvt-INs | yes | no |
| v7 | p7_m4 | 2 | 12500 | Hippo-dvt-INs | yes | no |
| v7 | p8_m3 | 0, 1, 6, 7, 9, | 112500 | Hippo-dvt-INs | yes | no |

| | | | | | | |
|---|---|---|---|---|---|---|
| | | 10, 11, 18, 24 | | | | |
| v7 | p8_m4 | 271 | 12500 | Hippo-dvt-INs | yes | no |
| v7 | p8_m5 | 0 | 12500 | Hippo-dvt-INs | yes | no |
| v7 | p11_m2 | 0, 2, 3 | 37500 | Hippo-dvt-INs | yes | no |
| v7 | p12_m1 | 0, 3, 7, 14 | 50000 | Hippo-dvt | yes | no |
| v7 | p6_m1 | 3 | 12500 | Hippo-dvt-INs | no | **8D** |
| v7 | p6_m2 | 10, 11, 13, 20 | 50000 | Hippo-dvt-INs | no | **8D** |
| v7 | p6_m3 | 11 | 12500 | Hippo-dvt-INs | no | **8D** |
| v7 | p8_m3 | 28, 32, 33 | 37500 | Hippo-dvt-INs | no | **8D** |
| v7 | p11_m2 | 4 | 12500 | Hippo-dvt-INs | no | **8D** |
